# A clinically validated human saliva metatranscriptomic test for global systems biology studies

**DOI:** 10.1101/2021.08.03.454950

**Authors:** Ryan Toma, Ying Cai, Oyetunji Ogundijo, Stephanie Gline, Diana Demusaj, Nathan Duval, Pedro Torres, Francine Camacho, Guruduth Banavar, Momchilo Vuyisich

**Affiliations:** Viome, Inc., Bothell and Bellevue, WA/New York, NY, USA, viome.com

**Author notes:** Corresponding author: Ryan Toma.

## Abstract

We report here the development of a high throughput, automated, inexpensive, and clinically validated saliva metatranscriptome test that requires less than 100 microliters of saliva. The RNA in the samples is preserved at the time of collection. Samples can then be stored and transported at ambient temperatures for up to 28 days without significantly affecting the test results. Another important feature of the sample preservative is that it inactivates all classes of pathogens, enabling safe transportation globally. Given the unique set of convenience, low cost, safety, and technical performance, this saliva metatranscriptomic test can be integrated into longitudinal, global scale, systems biology studies that will lead to an accelerated development of precision medicine diagnostic and therapeutic tools.

**Method Summary:** This report describes an important improvement to saliva transcriptome analysis. While current methods are complicated and expensive, the method reported here includes at-home sample collection, global shipping at ambient temperatures and pathogen inactivation at the point of collection; it uses fully automated, clinically validated and licensed laboratory and bioinformatic analyses.

## Introduction

Chronic diseases are the leading cause of morbidity and mortality globally. Yet, the majority of them have unknown etiologies, with weak genetic contributions. The human microbiome and alterations to human and microbial gene expression patterns have been identified as the underlying driver of many chronic diseases. To identify the etiology of chronic diseases and develop more effective preventative measures, a comprehensive gene expression analysis of the human body and associated microbiomes is needed, in addition to accurate strain-level taxonomy of all microorganisms. Saliva is easy to obtain and contains a diverse microbiome involved in many fundamental aspects of human physiology [1–3]. Saliva is also a suitable clinical specimen for the identification of human pathogens, such as SARS-CoV-2 [4]. Human transcripts in saliva can also produce an informative snapshot of human gene expression and their possible roles in human health and disease [5,6].

Human saliva contains a vast number of both commensal and potentially pathogenic microorganisms performing a wide variety of metabolic functions, with beneficial and potentially harmful effects to the host [7]. Human saliva also contains a substantial amount of human transcripts that can reveal specific interactions between the microbiome and human tissues [8]. Therefore, saliva presents an opportunity to investigate the oral microbiome and oral human transcriptome and their roles in disease pathogenesis [7,9]. In the era of precision medicine, methods to non-invasively assess an individual’s human and microbial gene expression levels can reveal predictive markers of disease, inform the best choice of therapy, and enable the development of novel treatment options. For example, using systems biology approaches to interrogate the transcriptome is now a standard strategy to stratify cancer patients and select the best available therapies [10–12]. Methods to analyze the transcriptome are becoming invaluable in the understanding of chronic diseases with unknown etiologies, but large scale adoption of metatranscriptomic analysis has not yet been possible due to the lack of low-cost and scalable methods.

The saliva microbiome has been associated with numerous chronic diseases including Alzheimer’s Disease [13], Parkinson’s Disease [14], Autism Spectrum Disorder [5], cancers [15], cardiovascular disease [16], diabetes [17], obesity [18], and autoimmune disorders [19,20]. As an example of the direct effect of salivary microorganisms on disease, *Fusobacterium nucleatum* originating in the oral cavity has been shown to promote colorectal cancer development [21,22]. The oral microbiome has also been shown to have a strong influence on the gut microbiome and the immune system, which are involved in a variety of different chronic diseases [23]. In addition to the clear role of the saliva microbiome in human health and disease, there is substantial information to be gained from human gene expression patterns in saliva. Alterations in salivary human gene expression and epigenetic markers have been observed in a range of disorders such as Autism Spectrum Disorder [6], Parkinson’s Disease [24], and Traumatic Brain Injury [25]. Saliva is clearly an informative sample type that can offer highly predictive insights into human health and disease and could provide information about the molecular mechanisms of disease onset and progression.

Despite the abundance of literature showing clear connections between the oral microbiome, human gene expression, and chronic diseases, most of the evidence is based on 16S ribosomal RNA gene sequencing or metagenomic (shotgun DNA) sequencing methods. These approaches limit the biomarker potential and actionability of interventions that can be made. 16S rRNA methods can provide taxonomic resolution to the genus level at best [26,27] and do not measure the biochemical functions of the microorganisms [28] or distinguish between living and dead microorganisms. In addition, traditional 16S rRNA gene sequencing excludes an analysis of human genes and excludes many bacteria and archaea, and all eukaryotic microorganisms and viruses [29], resulting in a limited view of the saliva microbiome ecosystem and minimizing the ability to discover novel microbial or human derived biomarkers.

Metagenomic sequencing can provide strain-level resolution of all DNA-based microorganisms [28], but it does not detect any RNA viruses (human, plant, and bacteriophages). Importantly, metagenomic analysis does not measure gene expression, it can only identify the *potential* biochemical functions of a microbial community. This is a disadvantage for studying disease states, which are affected more by gene expression and the biochemical outputs of the microbiome than the taxonomic identity of the microorganisms. This has already been demonstrated in several gut and oral microbiome studies [30,31]

The lack of comprehensive taxonomic classifications and the inability to investigate gene expression has limited the utility of the conventional methodologies (16S and metagenomics) in investigating the role of the saliva microbiome and human transcriptome in health and disease [32]. This is particularly apparent in the context of microbial functions, which have been shown to be more informative in disease states than taxonomy alone [30]. This showcases the utility of metatranscriptomic analysis of saliva, as it presents a more comprehensive and relevant snapshot of the microbial and human transcripts compared to traditional methodologies.

Saliva metatranscriptomic tests have already been used by our laboratory to create comprehensive and accurate diagnostic indicators of oral cancer [33]. Importantly, these predictive models rely on microbial taxonomy, microbial gene expression, and human gene expression signatures. Only through the use of a comprehensive sample analysis method, such as metatranscriptomics, can such a wide variety of data be analyzed.

In addition to their utility in biomarker discovery, functional outputs are also critical in developing a comprehensive understanding of disease etiology and subsequent treatment options [34]. Gene expression analyses can also provide information about specific microbial pathogenicity that is lost when simply looking at the microbiome composition. *Porphyromonas gingivalis* for example can be present in small amounts, but have highly expressed functions with a strong negative impact on the microbial community and ultimately the host [30,35].

Metatranscriptomic analyses address the limitations of both 16S rRNA gene sequencing and metagenomics. Metatranscriptomics provides insights into the biochemical activities of the microbiome by quantifying microbial gene expression levels, allowing for the assessment of biochemical pathway activities, while providing strain-level taxonomic resolution for all metabolically active microorganisms and viruses [36,37]. Metatranscriptomics also provides a comprehensive analysis of human genes in saliva. To date, metatranscriptomic methods have been limited due to the cost and complexity of both laboratory and bioinformatic methods [38]. Effective RNA preservation has been challenging, traditionally requiring a cold chain that is expensive and complicated. Another challenge of working with RNA is that about 97% of total RNA is ribosomal (rRNA) and 1% transfer RNA (tRNA), whereas messenger RNA (mRNA), which is the most informative RNA species, makes up the remaining 2%. By removing the non-informative prokaryotic and eukaryotic rRNAs and tRNA, more valuable transcriptome data can be generated with less sequencing depth [36,39], resulting in reduced per-sample sequencing costs and higher data resolution.

Here we present a comprehensive (sample collection-to-result) method for the quantitative metatranscriptomic analysis of saliva samples that can easily be applied to clinical studies and trials globally. The method is automated, high-throughput, inexpensive, clinically validated, and includes a fully automated bioinformatics suite for strain-level taxonomic classification of all microorganisms and human genes and their quantitative gene expression levels. An RNA preservation buffer mixed with the saliva sample at the point of collection inactivates all pathogens (bacteria, fungi, and viruses), preserves RNA integrity, enabling safe transportation globally at ambient temperatures for up to 28 days.

## Methods

### Ethics Statement

All procedures involving human subjects were performed in accordance with the ethical standards and approved by a federally-accredited Institutional Review Board (IRB) committee. Informed consent was obtained from all participants, who were residents of the USA at the time.

#### Sample collection

Saliva can be collected in any format that enables mixing of the Viome RNA preservative buffer (RPB) with the sample as soon as it is collected. Viome has developed a patented and convenient sample collection device that allows anyone, even children, to easily collect sufficient volumes of saliva and preserve it for metatranscriptomic analysis (Figure 1).

**Figure 1.**
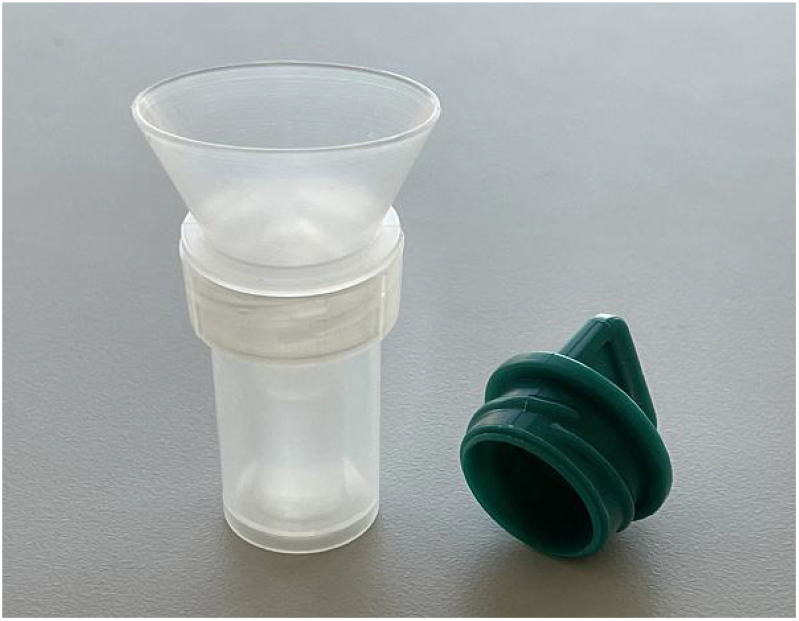
At-home saliva collection device. The insert contains the sample preservative that inactivates all types of pathogens, and preserves both RNA and DNA for 28 days at ambient temperature. Once saliva is deposited inside the tube, the funnel and insert are removed, which releases the preservative and mixes it with the specimen.

Participants were instructed to fast for 8 hours prior to collection, to collect the samples within one hour of waking up, and to gently rinse their mouth with water for ten seconds before collecting the sample.

### Method Validation

The performance of the saliva method was assessed by determining the method precision, sample stability, and longitudinal changes in saliva. Precision was assessed in four technical replicates of saliva from ten donors. Sample stability was assessed by storing samples for 7 and 28 days at room temperature (RT) compared to the technical replicates analyzed on the day of collection (day 0). A subset of replicate samples were shipped and returned to Viome to assess the impact of shipping on sample stability. All sample stability analyses were performed with four technical replicates between three donors. Longitudinal changes and helinger distances in saliva were assessed by collecting saliva weekly for five weeks from eight donors.

To develop a comprehensive saliva cohort, saliva samples were collected from 1,102 unique individuals. For the saliva cohort, the average age was 48 years (range: 12-88 years old) with 436 males, 665 females, and 1 participant identifying as another gender.

### Laboratory analysis

Saliva samples were lysed by bead beating in the RNA preservative buffer (RPB), as previously described [40,41]. RNA was extracted using silica beads and a series of washes, followed by elution in molecular biology grade water. DNA was degraded using RNase-free DNase. Prokaryotic and human rRNAs were removed using a subtractive hybridization method previously described [40]. Briefly, biotinylated DNA probes with sequences complementary to human and microbial rRNAs were added to total RNA, the mixture was heated and cooled, and the probe-rRNA complexes were removed using magnetic streptavidin beads. The remaining RNAs were converted to directional sequencing libraries with unique dual-barcoded adapters and ultrapure reagents. Libraries were pooled and quality controlled with dsDNA Qubit (Thermo Fisher Scientific) and Fragment Analyzer (Advanced Analytical) methods. Library pools were sequenced on Illumina NovaSeq instruments using 300 cycle kits.

### Bioinformatic analysis

Viome’s bioinformatics methods include quality control, strain-level taxonomic classification, microbial gene expression, and human gene expression characterizations. The quality control includes per-sample and per-batch metrics, such as the level of barcode hopping, batch contamination, positive and negative process controls, DNase efficacy, and number of reads obtained per sample. Following the quality control, the paired-end reads are aligned to a catalog containing ribosomal RNA (rRNA), human transcriptome, and 53,660 genomes spanning archaea, bacteria, fungi, protozoa, and viruses. Reads that map to rRNA are filtered out. Strain-level relative activities were computed from mapped reads via the expectation-maximization (EM) algorithm [42]. Relative activities at other levels of the taxonomic tree were then computed by aggregating according to taxonomic rank. Relative activities for the biological functions were computed by mapping paired-end reads to a catalog of 52,324,420 genes, quantifying gene-level relative activity with the EM algorithm and then aggregating gene-level activity by KEGG Ortholog (KO) annotation [43]. The identified and quantified active microbial species and KOs for each sample were then used for downstream analysis.

### Data analysis

Statistical parameters, including transformations and significance, are reported in the figures and figure legends. To compare pairs of samples, we report Spearman correlations (which are invariant to absolute expression levels of the genes and only consider the similarity of ranked expression) and Hellinger distance (an appropriate distance measure for compositional data). Multiplicative replacement method [44] is employed to deal with missing values, the data is transformed with centered log-ratio (CLR) [45] and Spearman correlation coefficients were computed. CLR transformation is commonly done to reduce false discoveries due to the compositional nature of sequencing data. Statistical analyses were performed in Python.

## Results

### Saliva Cohort Metrics

For the cohort of 1,102 saliva samples, the average number of sequencing reads per sample was 24,353,310. On average, there were 487 species, 2,539 microbial KOs, and 5,174 human transcripts detected per sample. See supplementary tables 1-5 for a rank ordered list of all detected genera, species, strains, microbial KOs, and human transcripts in the saliva cohort.

### Method precision

For longitudinal clinical studies in large populations that measure gene expression changes as a function of health and disease states, a method with high precision is extremely important, as it allows for the measurement of small changes in the levels of gene expression that are associated or predictive of chronic disease. To determine the precision of the metatranscriptomic method, Spearman correlations were calculated for microbial taxonomy (species) and microbial functions (KOs) for four technical replicates from ten participants (Figure 2a and 2b). For one participant (PID-0286), one replicate was determined to be an outlier with the log transformed effective sequencing depth (the number of sequencing reads aligned to microbial species) being greater than three standard deviations away from the mean and was removed from the analysis as it failed quality control. The method precision for microbial taxonomy and microbial functions is high, showing that the method is able to reliably reproduce the results (Figure 2a and 2b).

**Figure 2.**
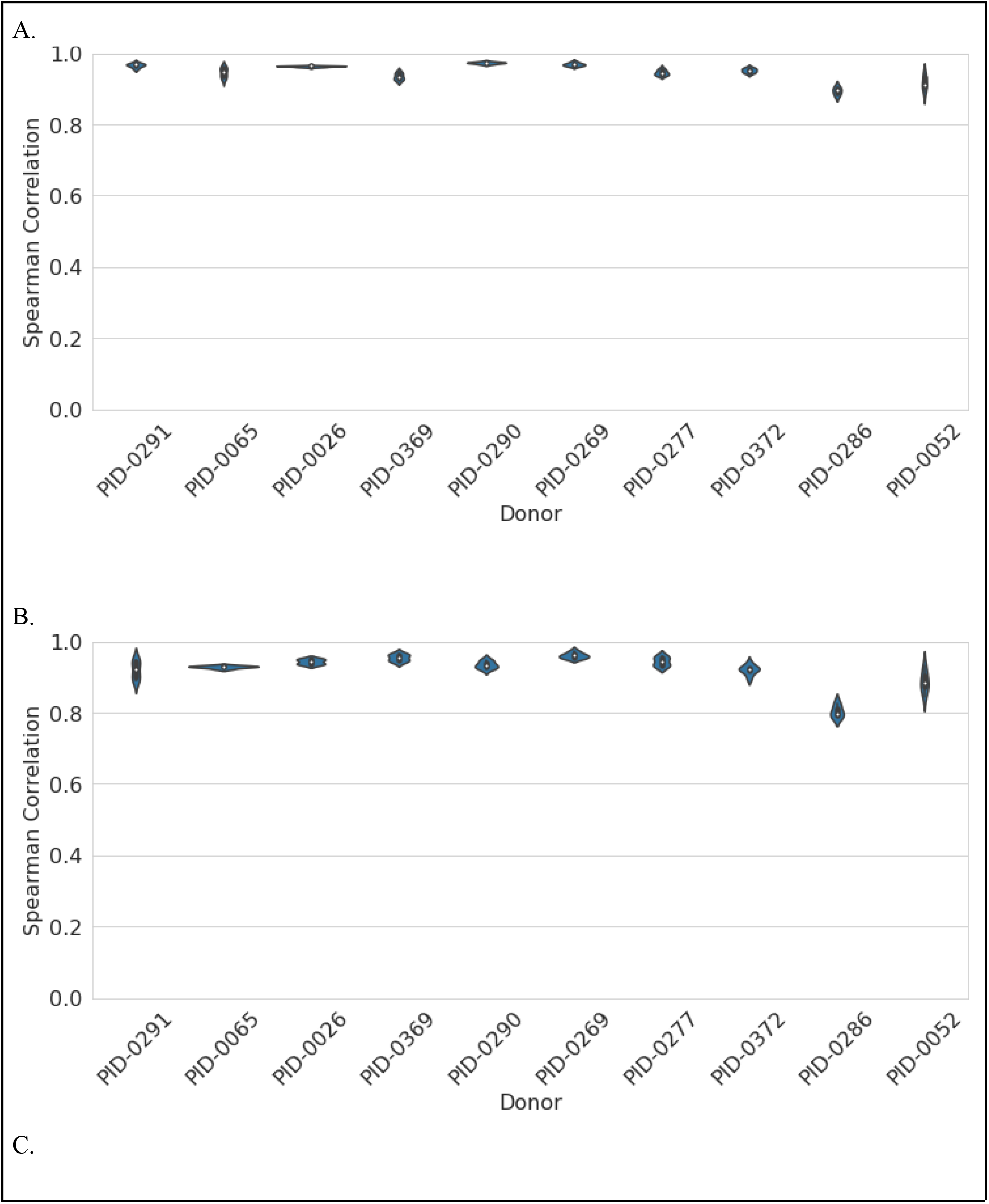
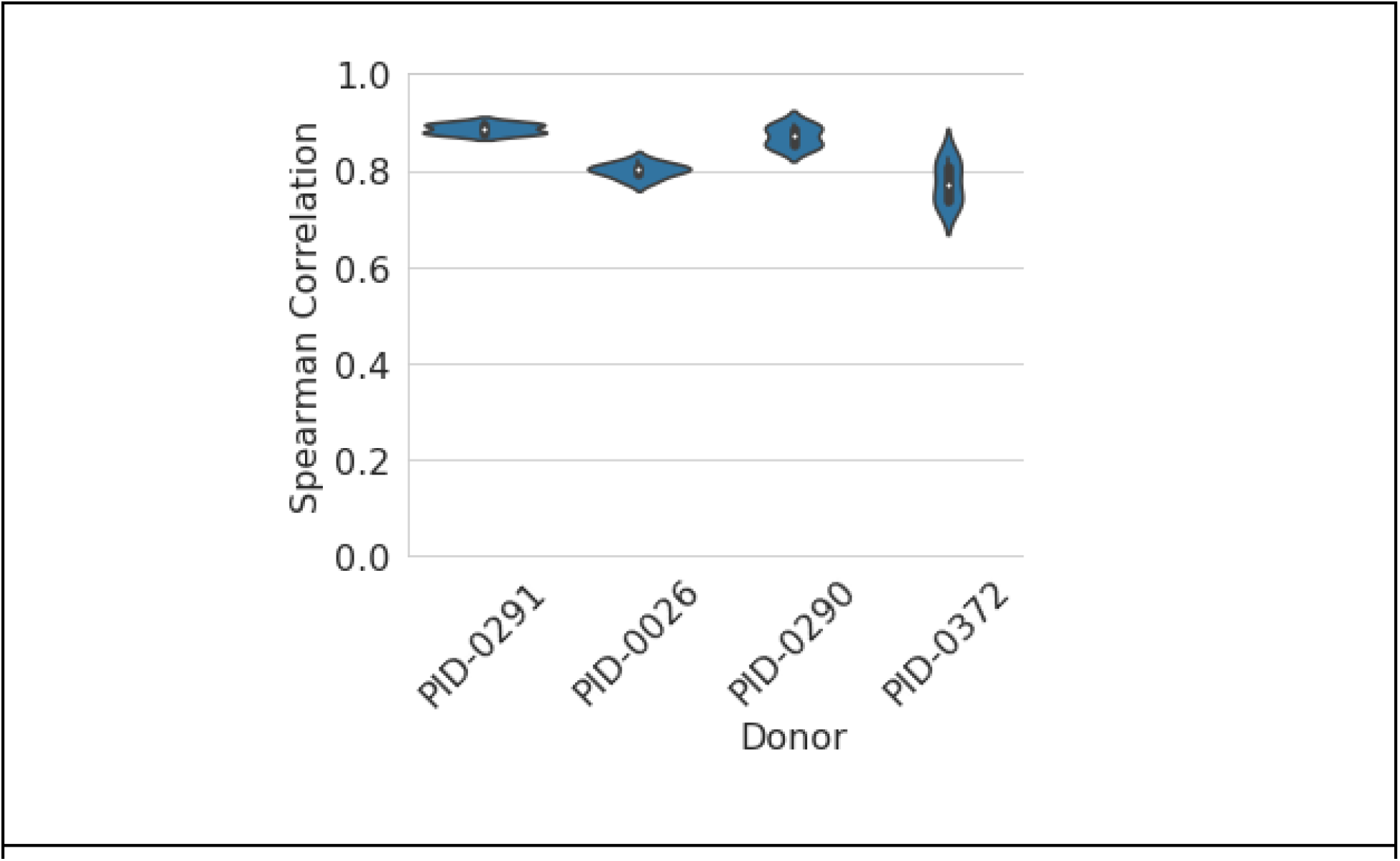
The method precision is shown comparing technical replicates from ten donors for microbial taxonomy (A), microbial functions, (B), and from four donors for human transcripts (C)

Human transcripts are present in low amounts in saliva and therefore require higher sequencing depth to analyze. Samples from four donors were re-sequenced with higher sequencing depth to determine the method precision for human genes. The effective sequencing depth (ESD, the number of reads aligned to human transcripts) needs to be about 1 million reads to yield high quality human gene expression data (table 1, Figure 2c). This demonstrates that the technology does allow for high precision human gene expression analyses from saliva, but requires higher sequencing depth.

**Table 1:**
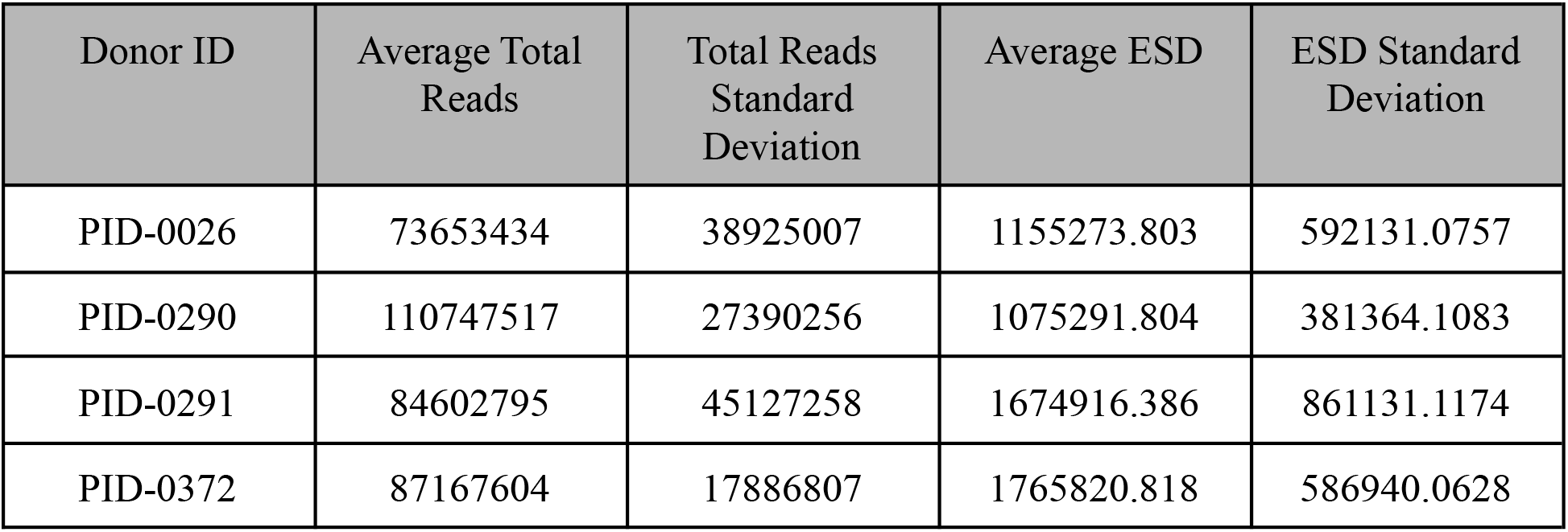
Total reads and ESD of re-sequenced saliva samples for human gene expression precision analyses.

| Donor ID | Average Total Reads | Total Reads Standard Deviation | Average ESD | ESD Standard Deviation |
| --- | --- | --- | --- | --- |
| PID-0026 | 73653434 | 38925007 | 1155273.803 | 592131.0757 |
| PID-0290 | 110747517 | 27390256 | 1075291.804 | 381364.1083 |
| PID-0291 | 84602795 | 45127258 | 1674916.386 | 861131.1174 |
| PID-0372 | 87167604 | 17886807 | 1765820.818 | 586940.0628 |

In addition to the Spearman correlations, two random technical replicates were compared from each participant for microbial taxonomy, microbial KOs, and human gene expression (Figure 3). These data show that Viome’s metatranscriptomic saliva test is able to produce data with high precision.

**Figure 3.**
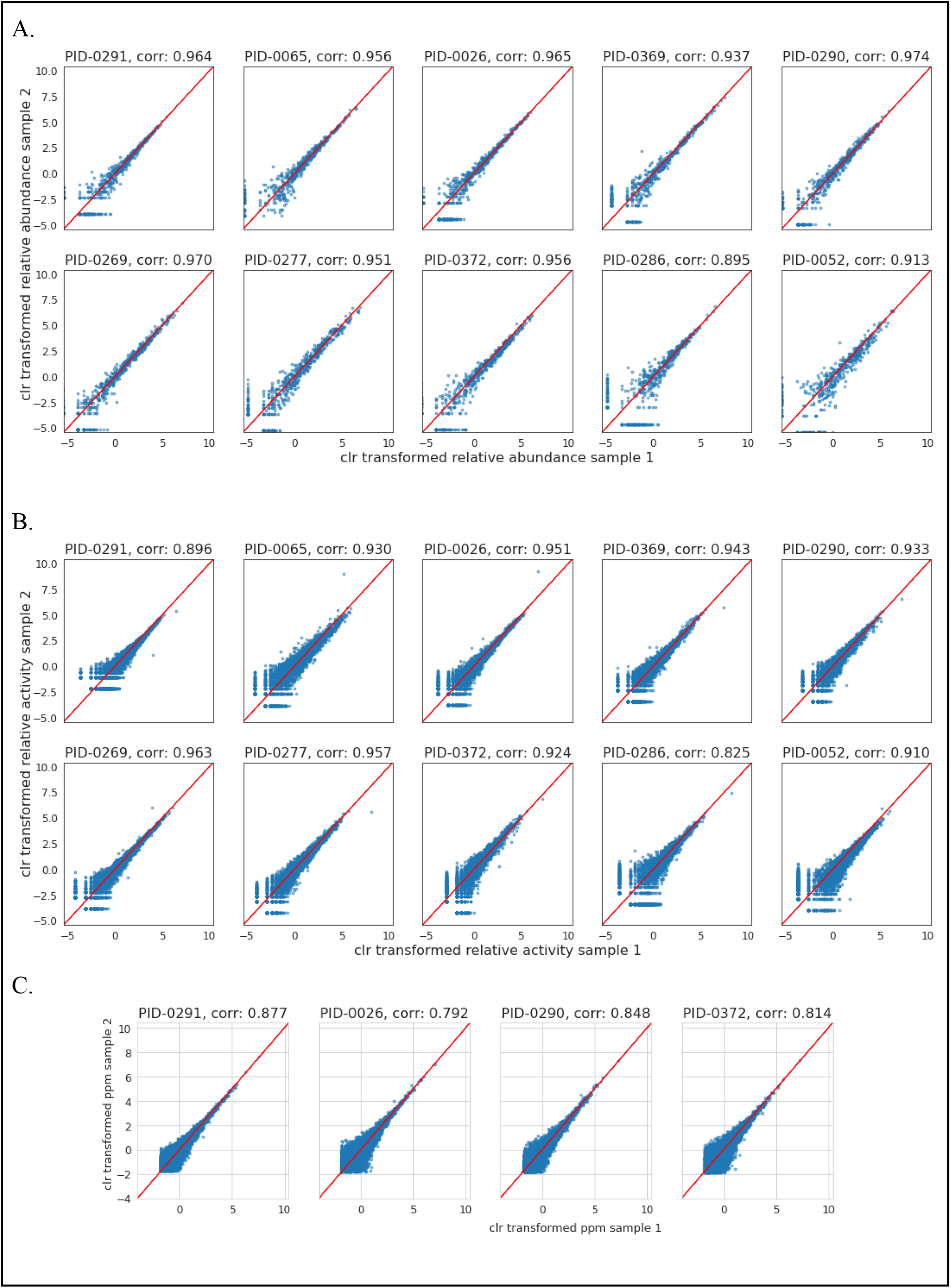
Relative abundance of taxons and transcripts are shown between two technical replicates from ten donors for microbial taxonomy (A), microbial functions (B), and from four donors for human genes (C)

### Sample stability

A major concern when performing metatranscriptomic analysis is the rapid and facile degradation of RNA, due to self-cleavage and action of ribonucleases [46]. To preserve RNA, our method uses an RNA preservative buffer (RPB) with multiple functions. RPB’s detergents quickly dissolve lipid bilayers and inactivate all microorganisms, the denaturants halt all enzymatic activity by disrupting the tertiary structures of proteins, while other ingredients prevent RNA self-cleavage by keeping the 2’ oxygen protonated. To validate the ability of RPB to preserve the saliva transcriptome at ambient temperatures, saliva samples were collected from three participants and stored at ambient temperature (72°F, 22°C) for 0, 7, and 28 days, with and without shipping. The shipping consisted of collecting the samples at the laboratory, sending them to an address via US Postal Service, then shipping them back to the laboratory using the same shipping method. Four technical replicates per condition were analyzed using our test, and microbial taxonomy and microbial functions were compared among all samples. The correlation between microbial species in saliva samples in all conditions is very high, with Spearman correlation coefficients above 0.928 for all conditions tested (Figure 4a). The correlation between microbial functions in saliva samples in all conditions is also very high, with Spearman correlation coefficients above 0.901 for all conditions tested (Figure 4b). This shows that Viome’s saliva metatranscriptomic analysis method can adequately preserve RNA for up to 28 days at ambient temperature, inclusive of sample shipping.

**Figure 4.**
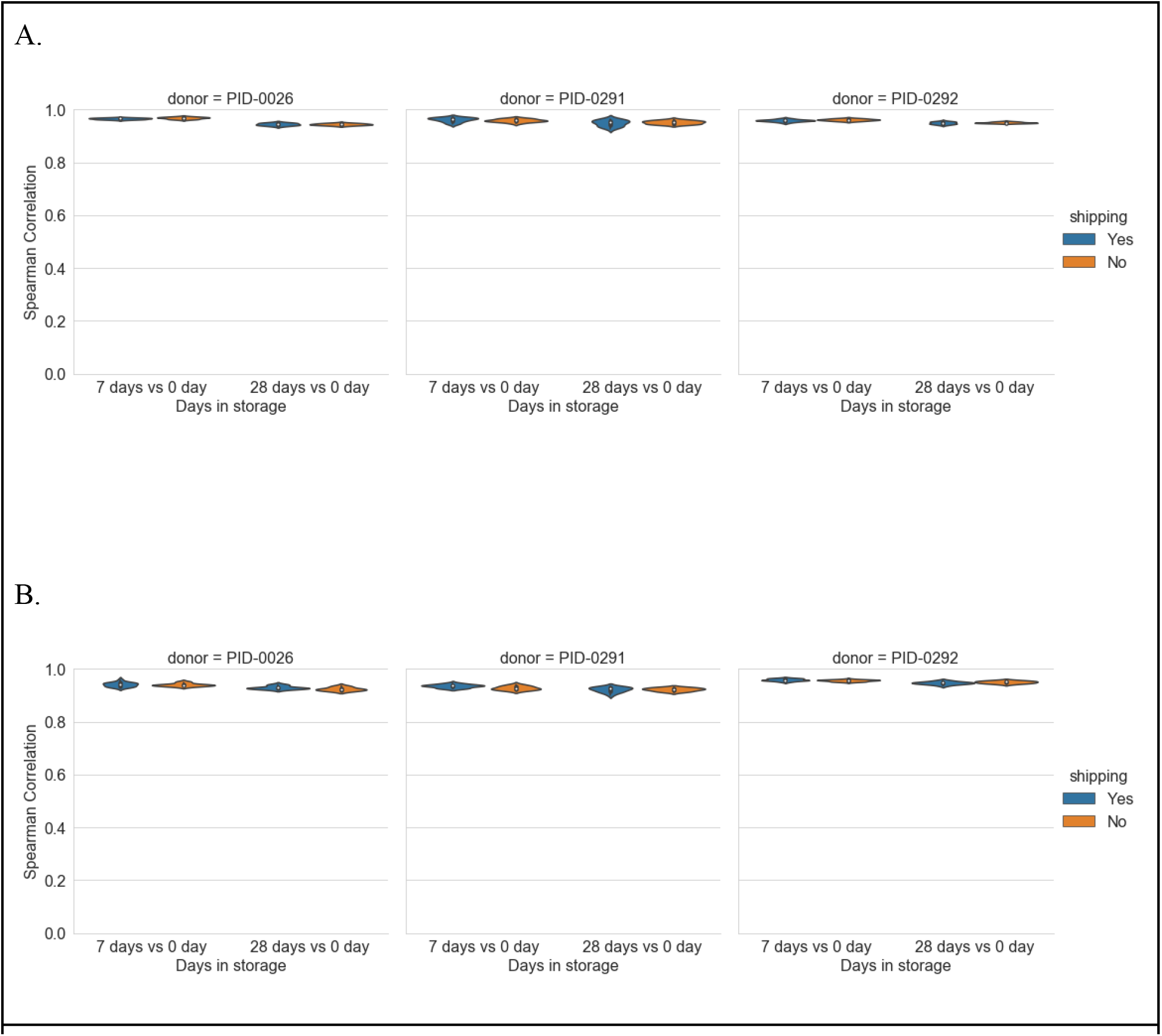
Spearman correlation coefficients of taxonomy (4a), and microbial functions (4b) for sample stability of saliva samples stored at ambient temperature for 0, 7, and 28 days, with and without shipping conditions.

### Longitudinal changes in the saliva transcriptome

It is important to understand the longitudinal stability of the salivary transcriptome so that large scale studies can be reliably carried out. Towards this goal, we recruited eight subjects that collected saliva samples weekly for five weeks. On average, 537.18 microbial species and 2693.54 microbial KOs were detected per sample. For all of the taxa and KOs that were detected, the correlation remains very high across five weeks for each participant (Figure 5 and Supplemental Figure 1). The correlations observed between the 1 week and 5 week samples for taxonomy ranged from 0.87 to 0.94 (Figure 5A and Supplemental Figure 1 A-H) and for KOs ranged from 0.88 to 0.94 (Figure 5B and Supplemental Figure 1 I-P). These data show that the saliva microbiome is longitudinally stable both in terms of composition (taxonomy) and activity (KOs).

**Figure 5.**
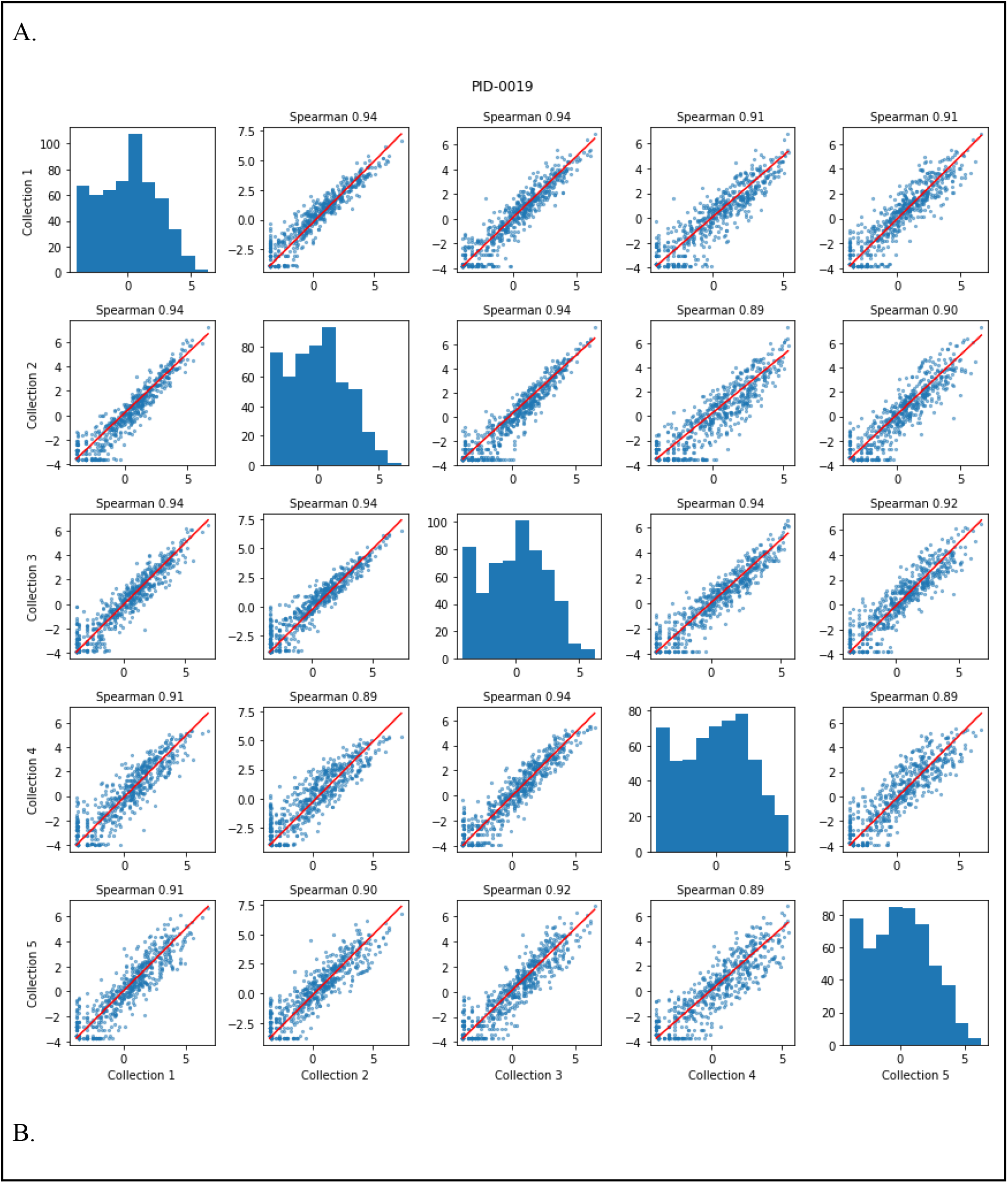
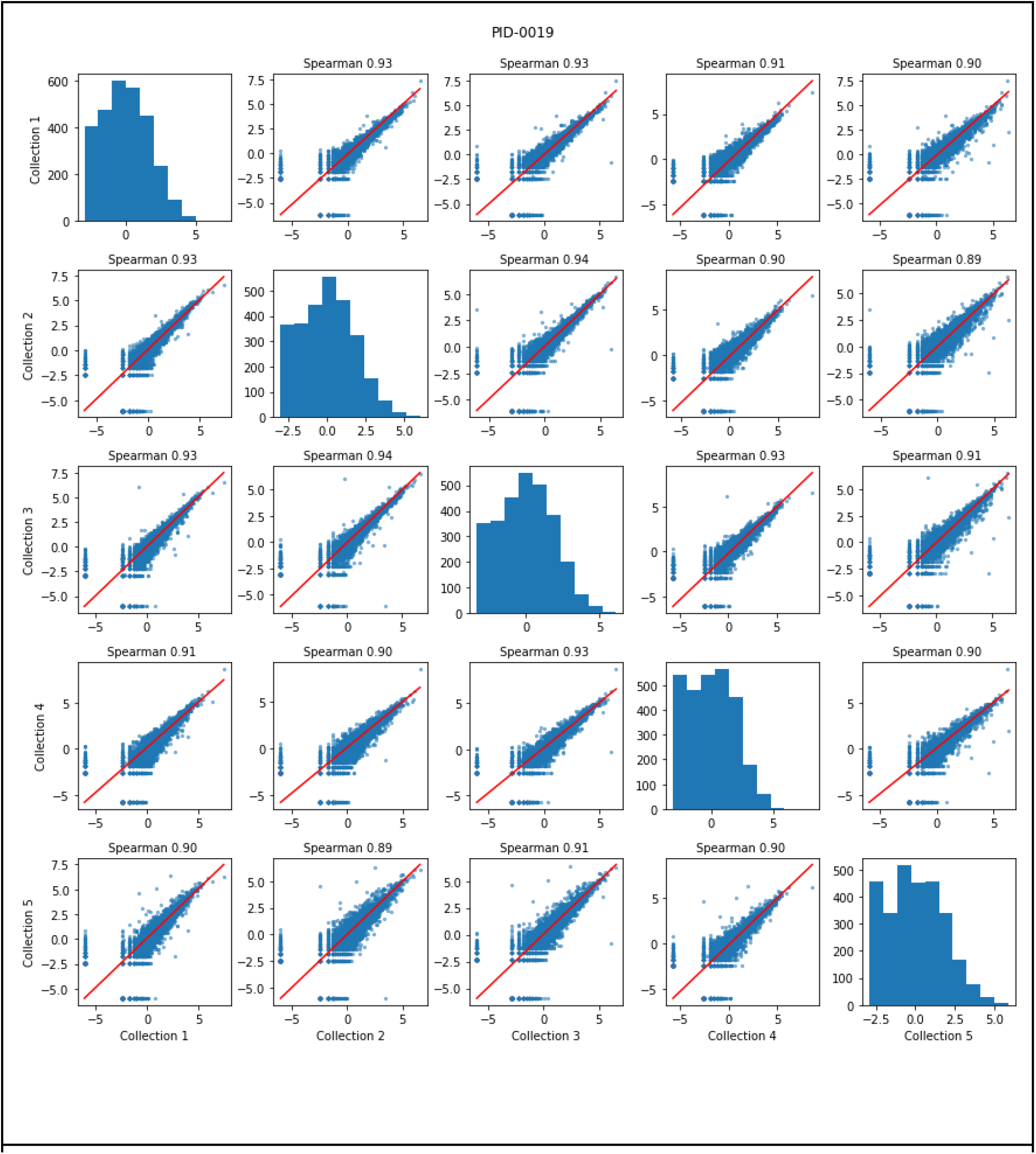
Longitudinal stability of saliva transcriptome in eight study participants over a 5 week period (one participant is shown here, for all eight participants see Supplementary Figure 1). (A) Scatter plots comparing the relative abundance values of each taxon at each collection time. (B) The same analysis was repeated for KOs comparing sum transcripts per million (tpm) of each KO.

### Method Precision

One of the more important parameters of the saliva test is to be sufficiently precise to distinguish very small changes among thousands of measured features. Such high precision would enable the identification of transcriptome changes over time, and changes related to health and disease. To assess the test precision in this context, we compared Hellinger distances among Biological samples from the same person collected over time (intraperson distances in Figure 6), and Biological samples obtained from different people (interperson distances in Figure 6). Empirical cumulative distribution function (ECDF) plots of Hellinger distance show that when taking microbial species (Figure 6A), microbial functions (Figure 6B), and human genes (Figure 6C) into account, samples taken from one individual over five different weeks (intraperson distances) tend to be more similar than pairs of samples coming from two different participants (interperson distances).

**Figure 6.**
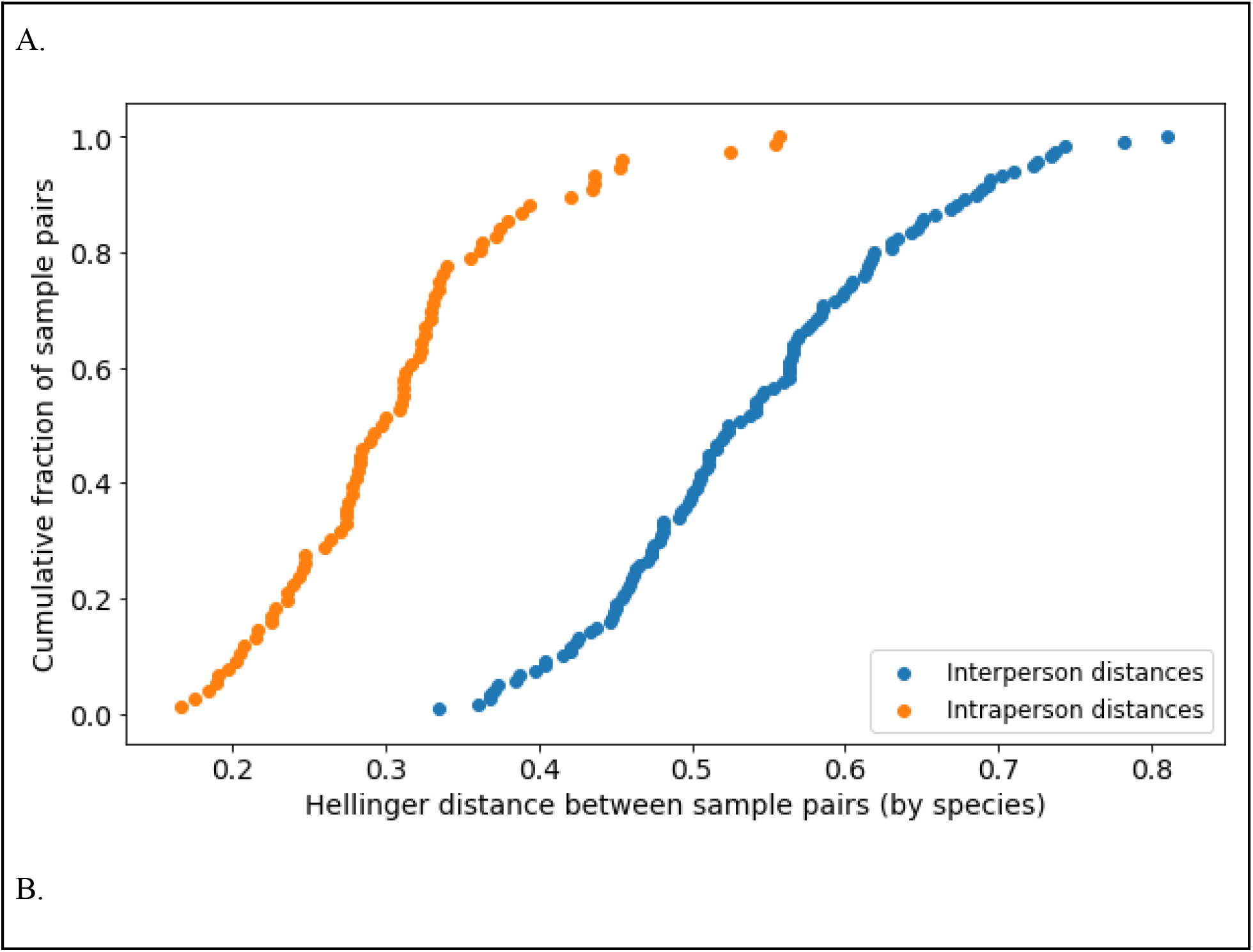
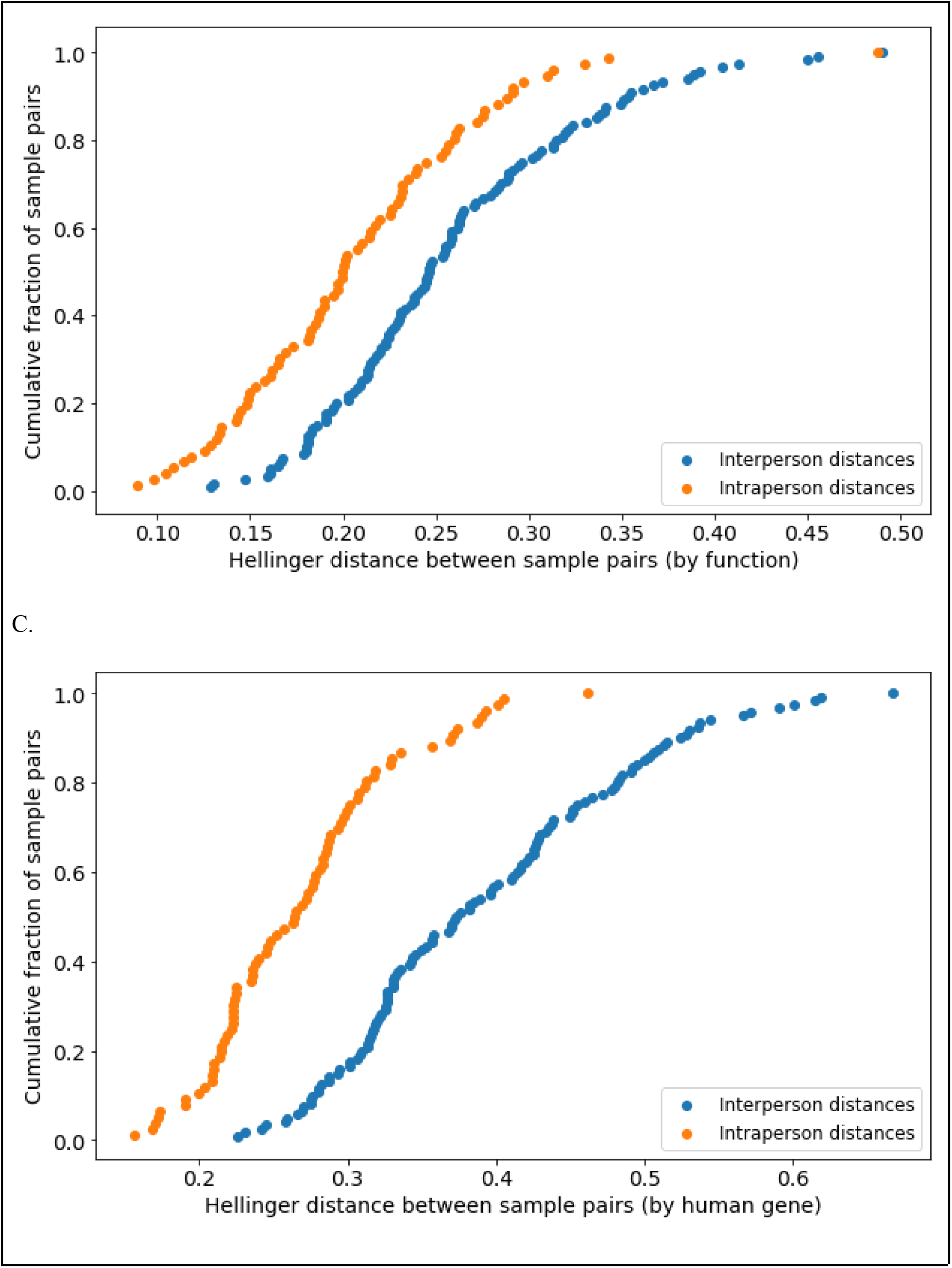
Empirical cumulative distribution function (ECDF) plots of Hellinger distance shows that when taking microbial species (A), microbial functions (B), and human genes (C) into account, samples taken from one individual over five different weeks (intraperson distances) tend to be more similar than pairs of samples coming from two different participants (interperson distances).

## Discussion

Chronic, non-communicable diseases are the leading cause of death globally [47]. There is strong evidence that human health, aging, and chronic diseases are heavily influenced by nutrition and microbiome functions. To understand the root causes of chronic diseases, we need to study the human body as an ecosystem, which requires a systems biology approach. Gene expression levels, for both the human and microbiome, play a very important role in chronic disease onset and progression, and need to be studied in the context of an integrated systems biology platform. While scalable and clinically validated gut microbiome and blood transcriptome gene expression tests have been developed [41,48], a human saliva transcriptome test with the same features and performance is also needed.

Here we describe a novel saliva transcriptome test that is clinically validated and globally scalable. This is the first saliva test capable of producing high quality microbial taxonomic classifications and both microbial and human gene expression profiles. The saliva sample (87.5 microliters) can be collected by anyone, anywhere, and can be stored and shipped at ambient temperature to a reference laboratory. The method is automated, inexpensive, and high throughput. The included preservative (RPB) inactivates pathogens, which is very important for international shipping, as it prevents disease spread. RPB also preserves RNA for up to 28 days at ambient temperature, which obviates the need for cold storage and specialized shipping, which are expensive and not easily available. The method precision is high and sufficient to study populations.

## Future Directions

This test, along with the previously published stool and blood metatranscriptome tests, can be integrated into a comprehensive systems biology platform that can be applied to population-scale longitudinal studies. Such large studies will identify mechanisms of chronic disease onset and progression, improve or bring new diagnostic and companion diagnostic tools, and enable precision nutrition and microbiome engineering to prevent and cure chronic diseases.

## Author Contributions

M Vuyisich and R Toma conceived and designed the methods. R Toma, D Demusaj, and N Duval performed data collection. Y Cai, O Ogundijo, S Gline, and G Banavar performed data analysis. All authors contributed to data interpretation. R Toma and N Duval contributed to the writing of the manuscript.

## Ethical conduct of research

The authors state that they have obtained appropriate institutional review board approval or have followed the principles outlined in the Declaration of Helsinki for all human or animal experimental investigations. In addition, for investigations involving human subjects, informed consent has been obtained from the participants involved.

## Financial and competing interests disclosure

All authors are current or former employees of Viome, Inc. The authors have no other relevant affiliations or financial involvement with any organization or entity with a financial interest in or financial conflict with the subject matter or materials discussed in the manuscript apart from those disclosed.

No writing assistance was utilized in the production of this manuscript.

## Notes

### Summary of Updates

Pedro Torres and Francine Camacho added as authors

