## Supplemental Figure 1 for "A clinically validated human saliva metatranscriptomic test for global systems biology studies"

A.

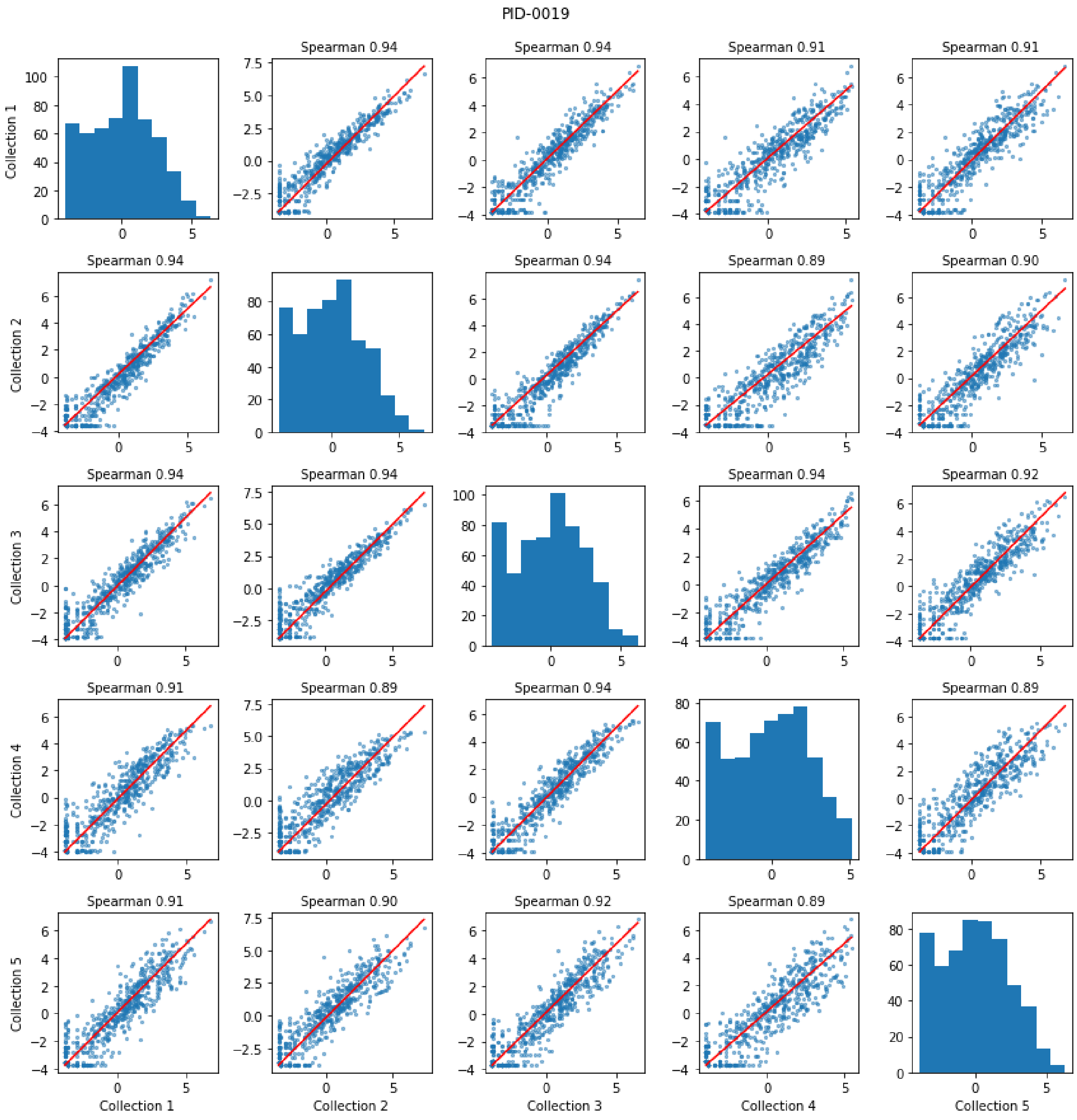
PID-0019
Spearman 0.94
Spearman 0.94
Spearman 0.91
Spearman 0.91
Spearman 0.94
Spearman 0.94
Spearman 0.89
Spearman 0.90
Spearman 0.94
Spearman 0.94
Spearman 0.94
Spearman 0.92
Spearman 0.91
Spearman 0.89
Spearman 0.94
Spearman 0.89
Spearman 0.91
Spearman 0.90
Spearman 0.92
Spearman 0.89
Collection 1
Collection 2
Collection 3
Collection 4
Collection 5

B.

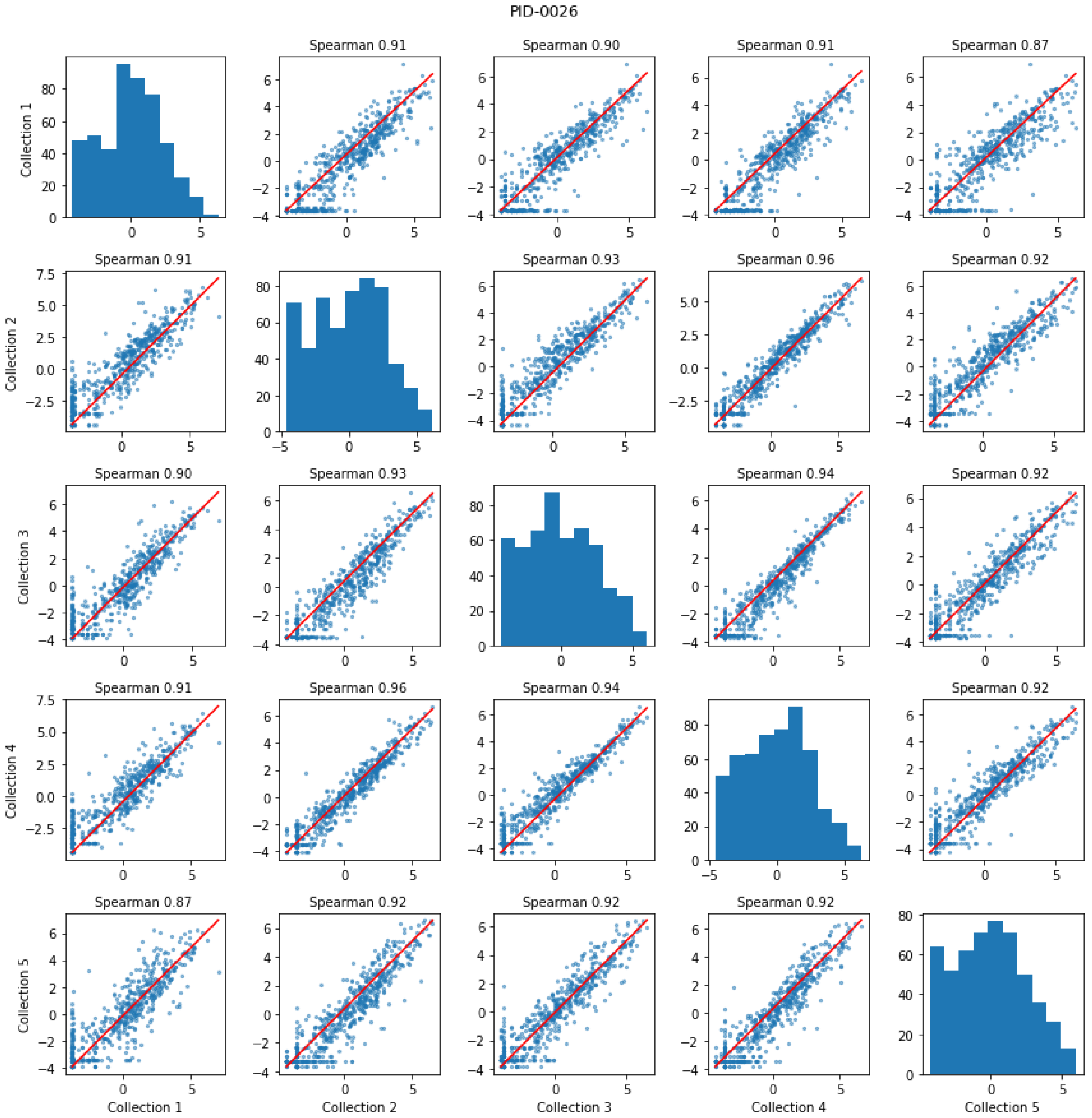
PID-0026
Spearman 0.91
Spearman 0.90
Spearman 0.91
Spearman 0.87
Spearman 0.91
Spearman 0.93
Spearman 0.96
Spearman 0.92
Spearman 0.90
Spearman 0.93
Spearman 0.94
Spearman 0.92
Spearman 0.91
Spearman 0.96
Spearman 0.94
Spearman 0.92
Spearman 0.87
Spearman 0.92
Spearman 0.92
Spearman 0.92
Collection 1
Collection 2
Collection 3
Collection 4
Collection 5

C.

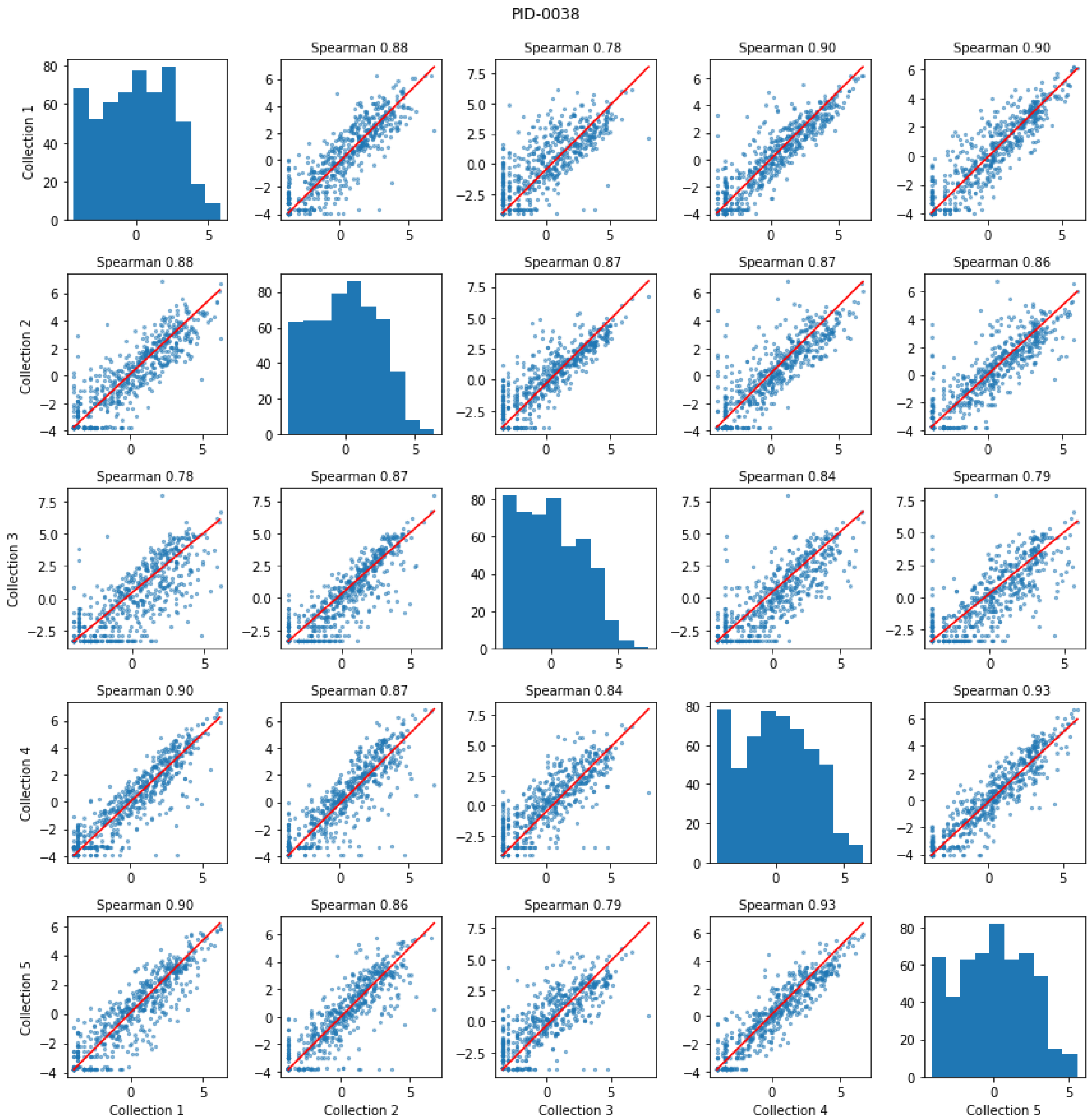
PID-0038
Spearman 0.88
Spearman 0.78
Spearman 0.90
Spearman 0.90
Spearman 0.88
Spearman 0.87
Spearman 0.87
Spearman 0.86
Spearman 0.78
Spearman 0.87
Spearman 0.84
Spearman 0.79
Spearman 0.90
Spearman 0.87
Spearman 0.84
Spearman 0.93
Spearman 0.90
Spearman 0.86
Spearman 0.79
Spearman 0.93
Collection 1
Collection 2
Collection 3
Collection 4
Collection 5

D.

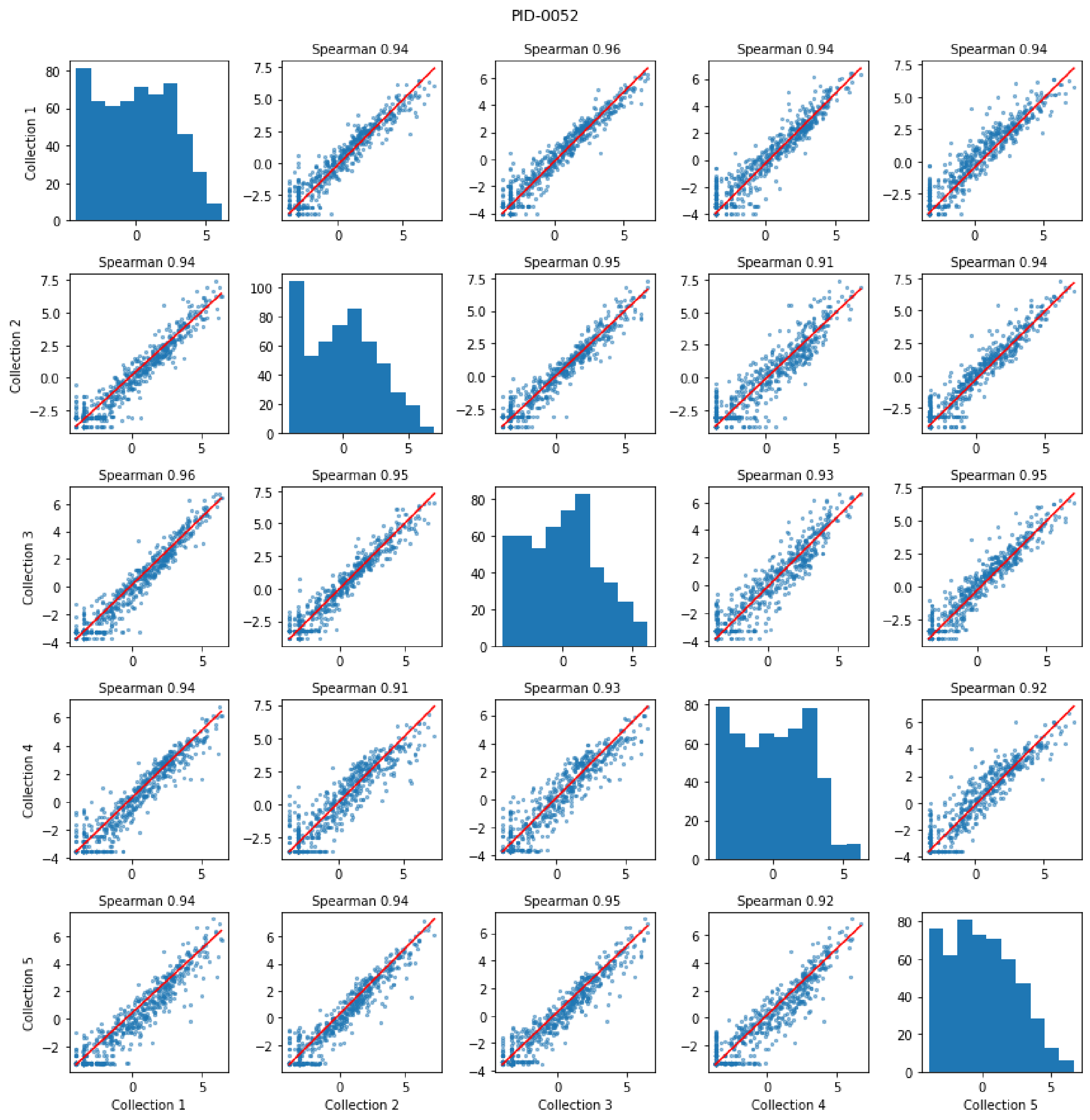
PID-0052
Spearman 0.94
Spearman 0.96
Spearman 0.94
Spearman 0.94
Spearman 0.94
Spearman 0.95
Spearman 0.91
Spearman 0.94
Spearman 0.96
Spearman 0.95
Spearman 0.93
Spearman 0.95
Spearman 0.94
Spearman 0.91
Spearman 0.93
Spearman 0.92
Spearman 0.94
Spearman 0.94
Spearman 0.95
Spearman 0.92
Collection 1
Collection 2
Collection 3
Collection 4
Collection 5

E.

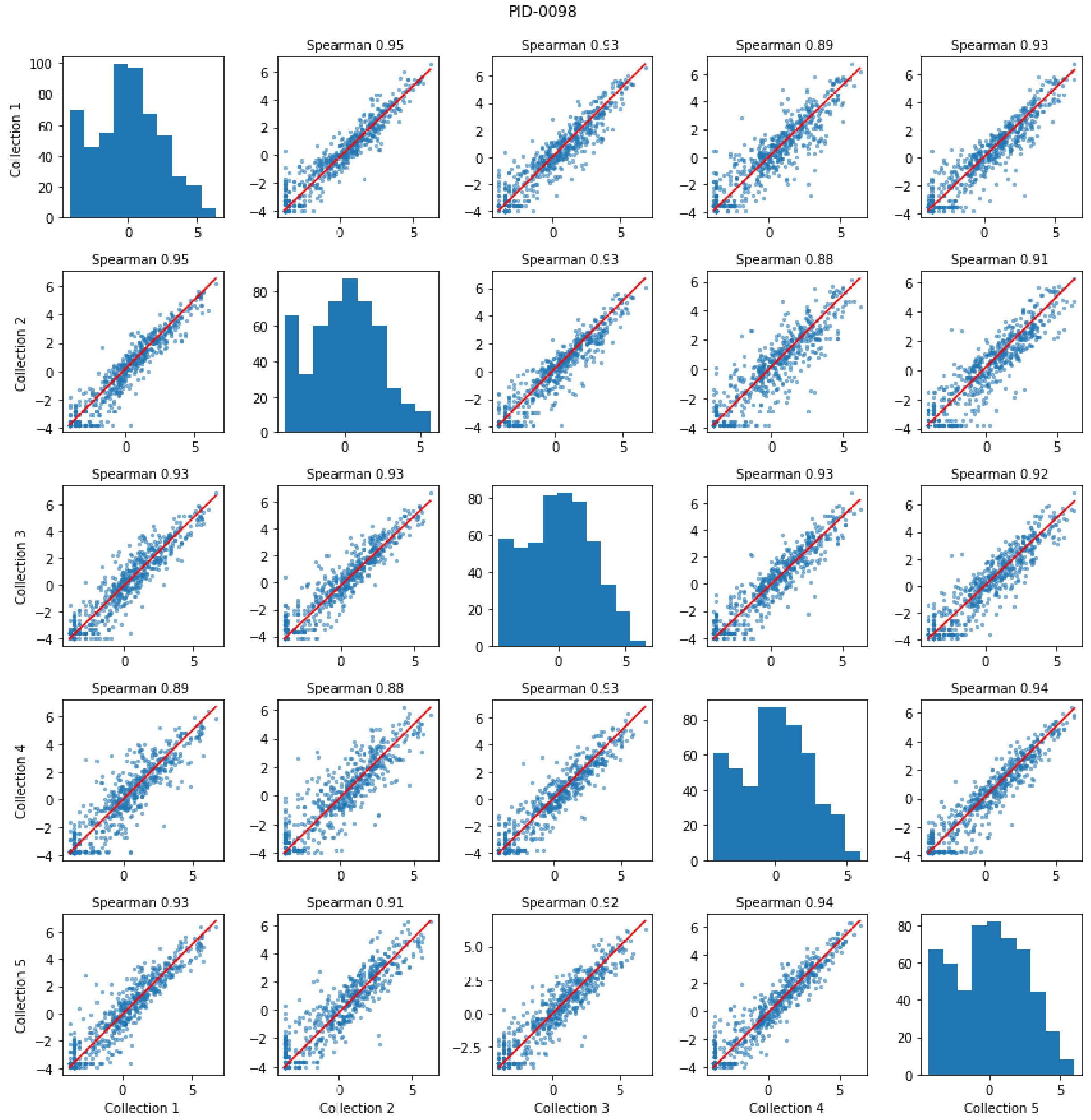
PID-0098
Spearman 0.95
Spearman 0.93
Spearman 0.89
Spearman 0.93
Spearman 0.95
Spearman 0.93
Spearman 0.88
Spearman 0.91
Spearman 0.93
Spearman 0.93
Spearman 0.93
Spearman 0.92
Spearman 0.89
Spearman 0.88
Spearman 0.93
Spearman 0.94
Spearman 0.93
Spearman 0.91
Spearman 0.92
Spearman 0.94
Collection 1
Collection 2
Collection 3
Collection 4
Collection 5

F.

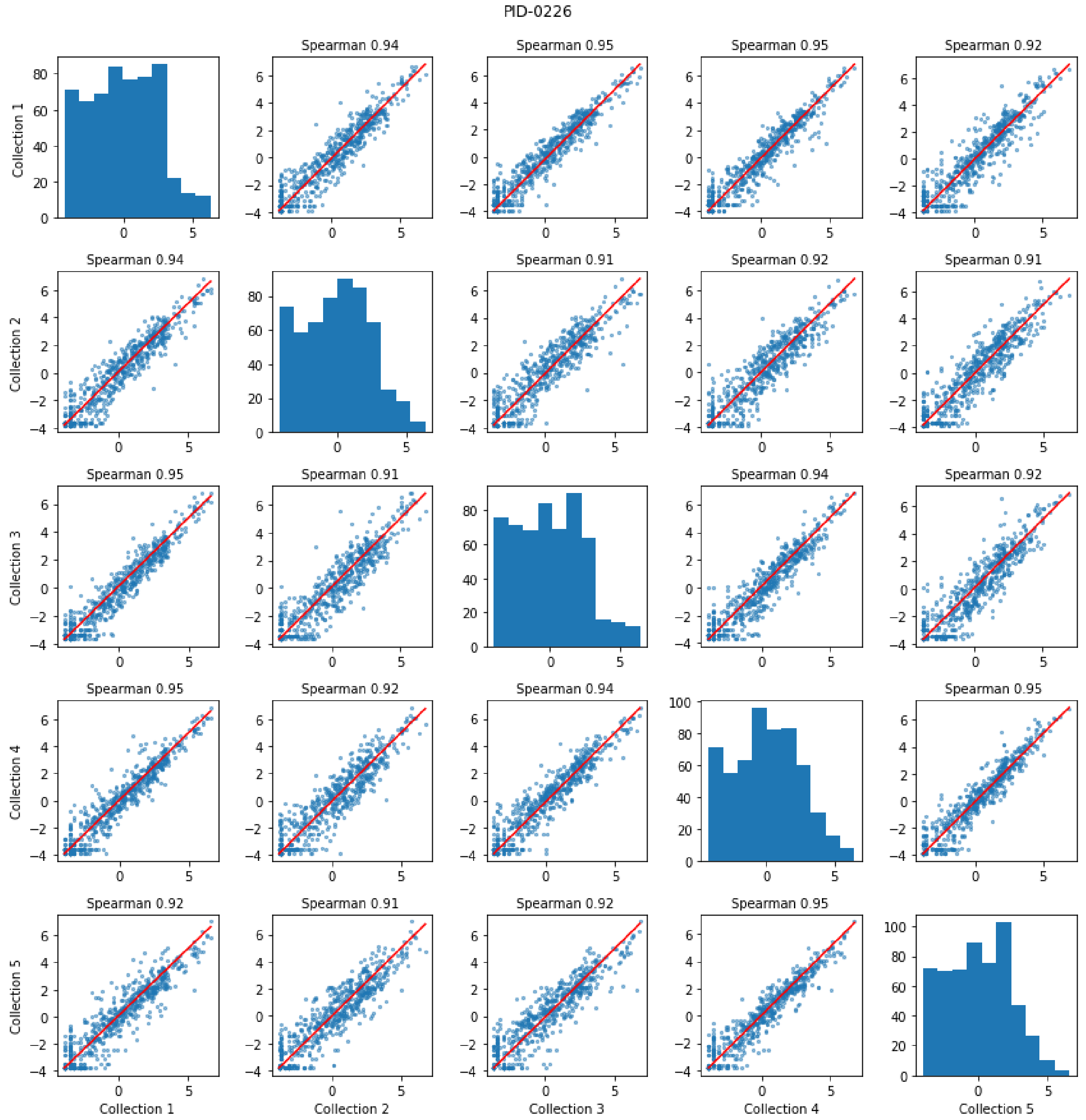
PID-0226
Spearman 0.94
Spearman 0.95
Spearman 0.95
Spearman 0.92
Spearman 0.94
Spearman 0.91
Spearman 0.92
Spearman 0.91
Spearman 0.95
Spearman 0.91
Spearman 0.94
Spearman 0.92
Spearman 0.95
Spearman 0.92
Spearman 0.94
Spearman 0.95
Spearman 0.92
Spearman 0.91
Spearman 0.92
Spearman 0.95
Collection 1
Collection 2
Collection 3
Collection 4
Collection 5

G.

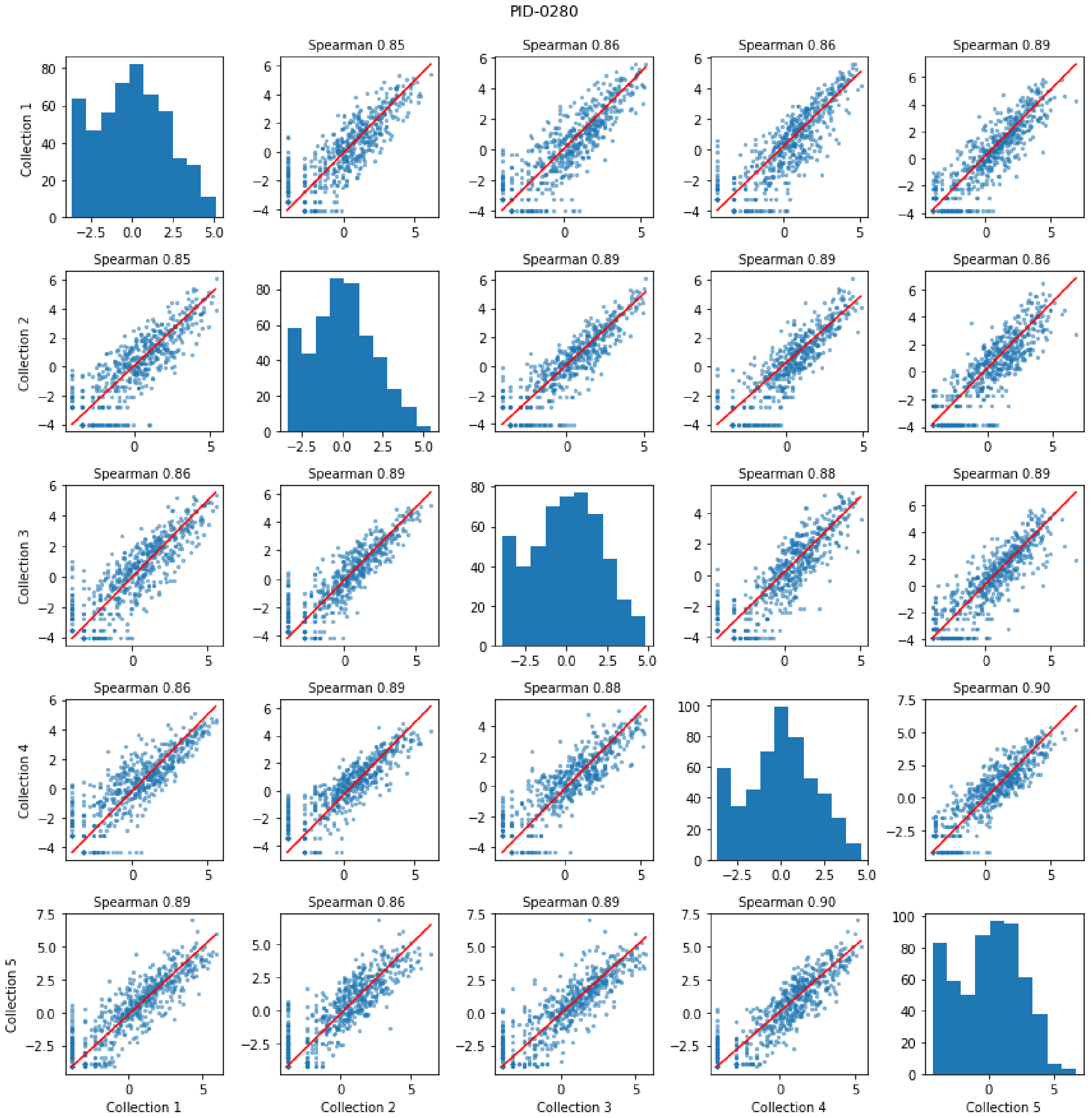
PID-0280
Spearman 0.85
Spearman 0.86
Spearman 0.86
Spearman 0.89
Spearman 0.85
Spearman 0.89
Spearman 0.89
Spearman 0.86
Spearman 0.86
Spearman 0.89
Spearman 0.88
Spearman 0.89
Spearman 0.86
Spearman 0.89
Spearman 0.88
Spearman 0.90
Spearman 0.89
Spearman 0.86
Spearman 0.89
Spearman 0.90
Collection 1
Collection 2
Collection 3
Collection 4
Collection 5

H.

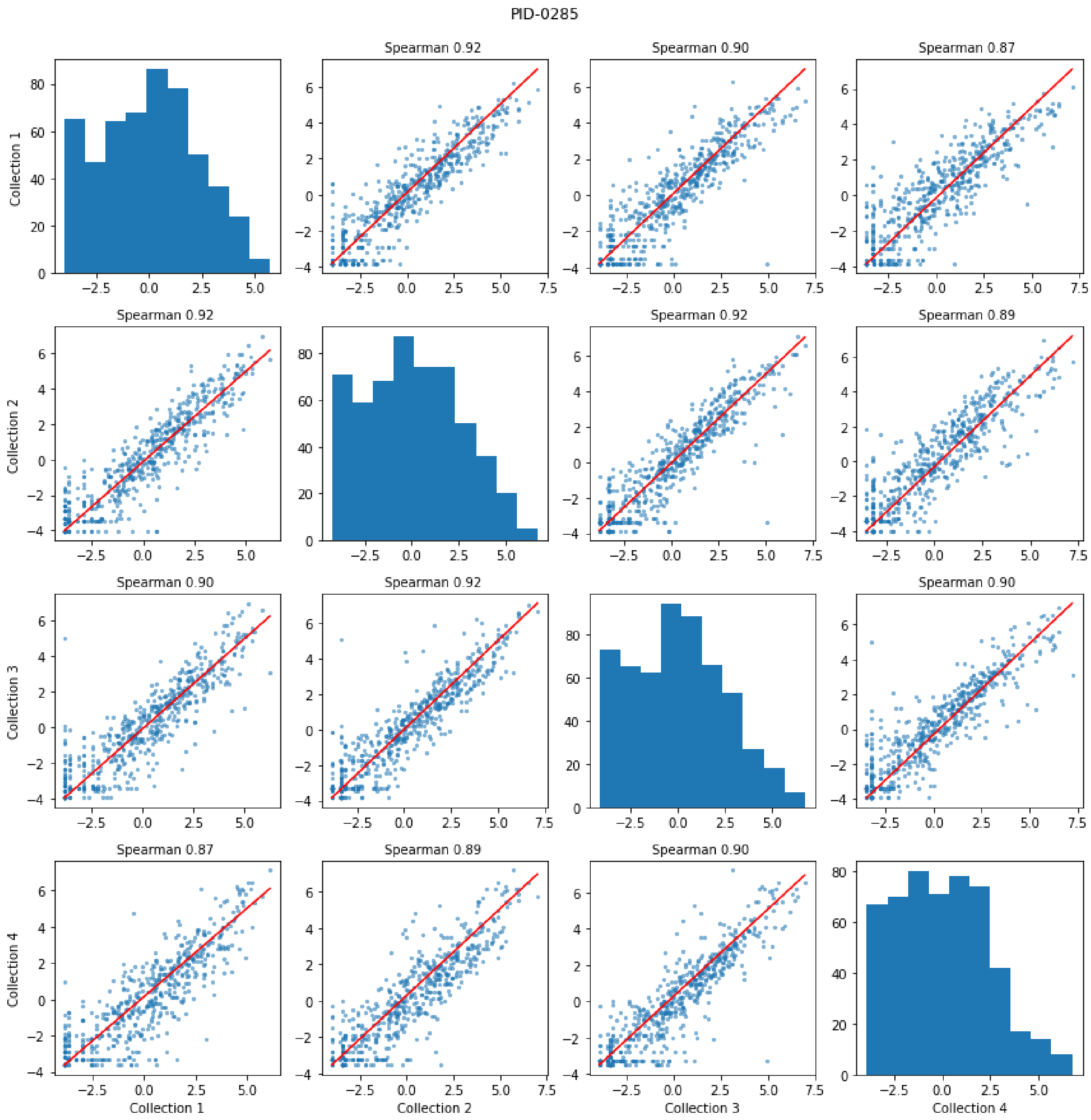
PID-0285
Spearman 0.92
Spearman 0.90
Spearman 0.87
Spearman 0.92
Spearman 0.92
Spearman 0.89
Spearman 0.90
Spearman 0.92
Spearman 0.90
Spearman 0.87
Spearman 0.89
Spearman 0.90
Collection 1
Collection 2
Collection 3
Collection 4

I.

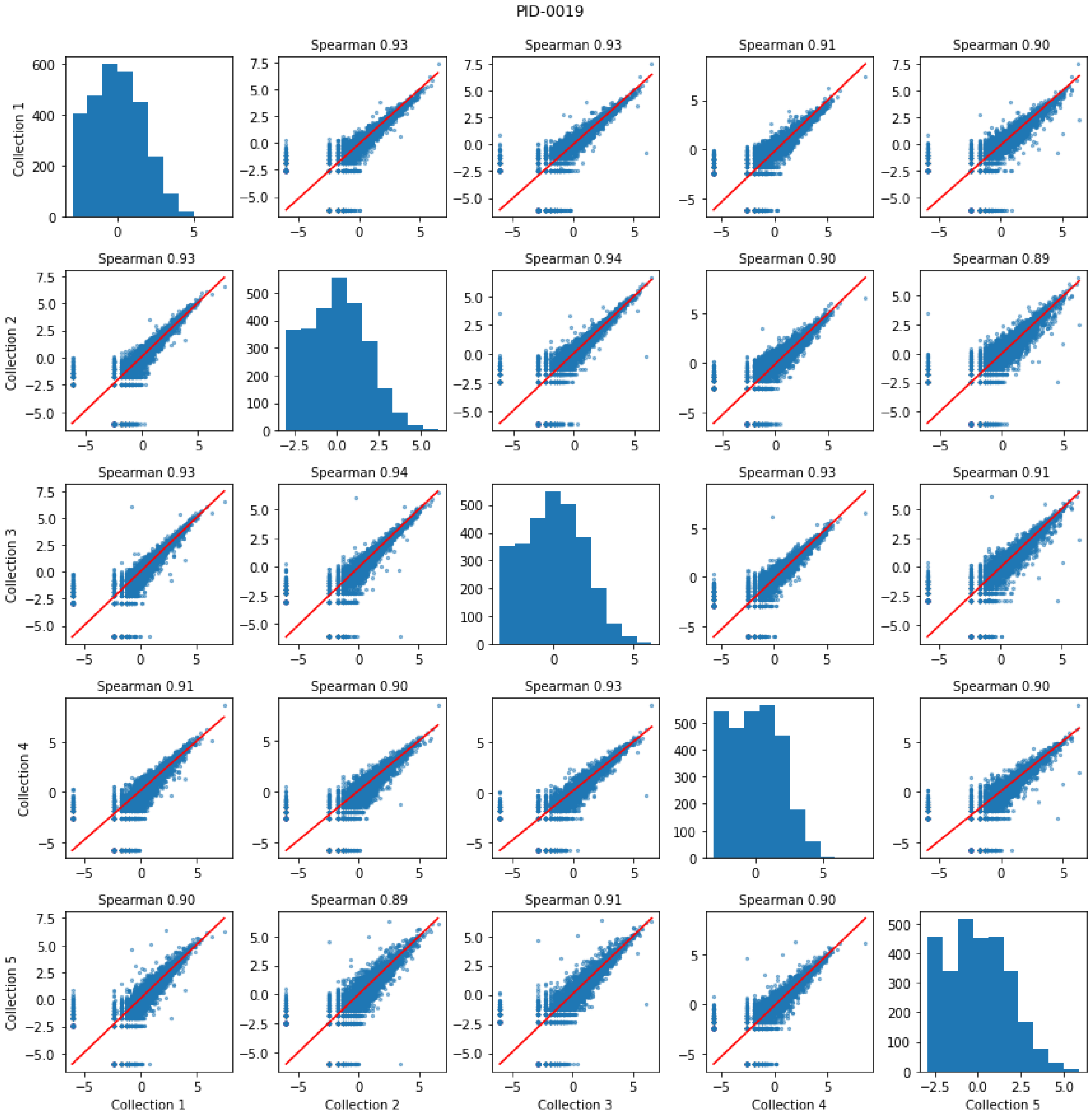

PID-0019
Spearman 0.93
Spearman 0.93
Spearman 0.91
Spearman 0.90
Spearman 0.93
Spearman 0.94
Spearman 0.90
Spearman 0.89
Spearman 0.93
Spearman 0.94
Spearman 0.93
Spearman 0.91
Spearman 0.91
Spearman 0.90
Spearman 0.93
Spearman 0.90
Spearman 0.90
Spearman 0.89
Spearman 0.91
Spearman 0.90
Collection 1
Collection 2
Collection 3
Collection 4
Collection 5


J.

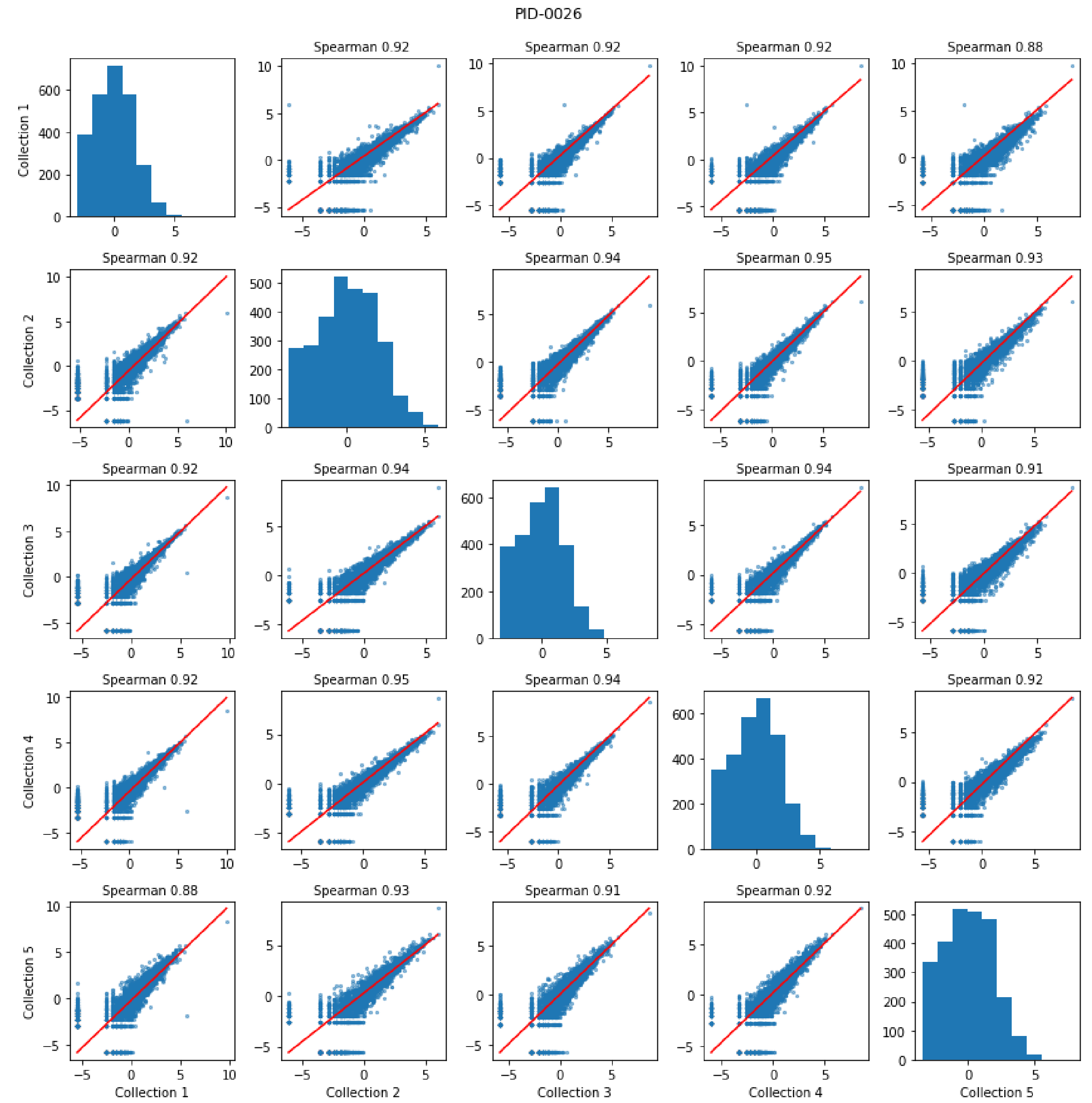
PID-0026
Spearman 0.92
Spearman 0.92
Spearman 0.92
Spearman 0.88
Spearman 0.92
Spearman 0.94
Spearman 0.95
Spearman 0.93
Spearman 0.92
Spearman 0.94
Spearman 0.94
Spearman 0.91
Spearman 0.92
Spearman 0.95
Spearman 0.94
Spearman 0.92
Spearman 0.88
Spearman 0.93
Spearman 0.91
Spearman 0.92
Collection 1
Collection 2
Collection 3
Collection 4
Collection 5

K.

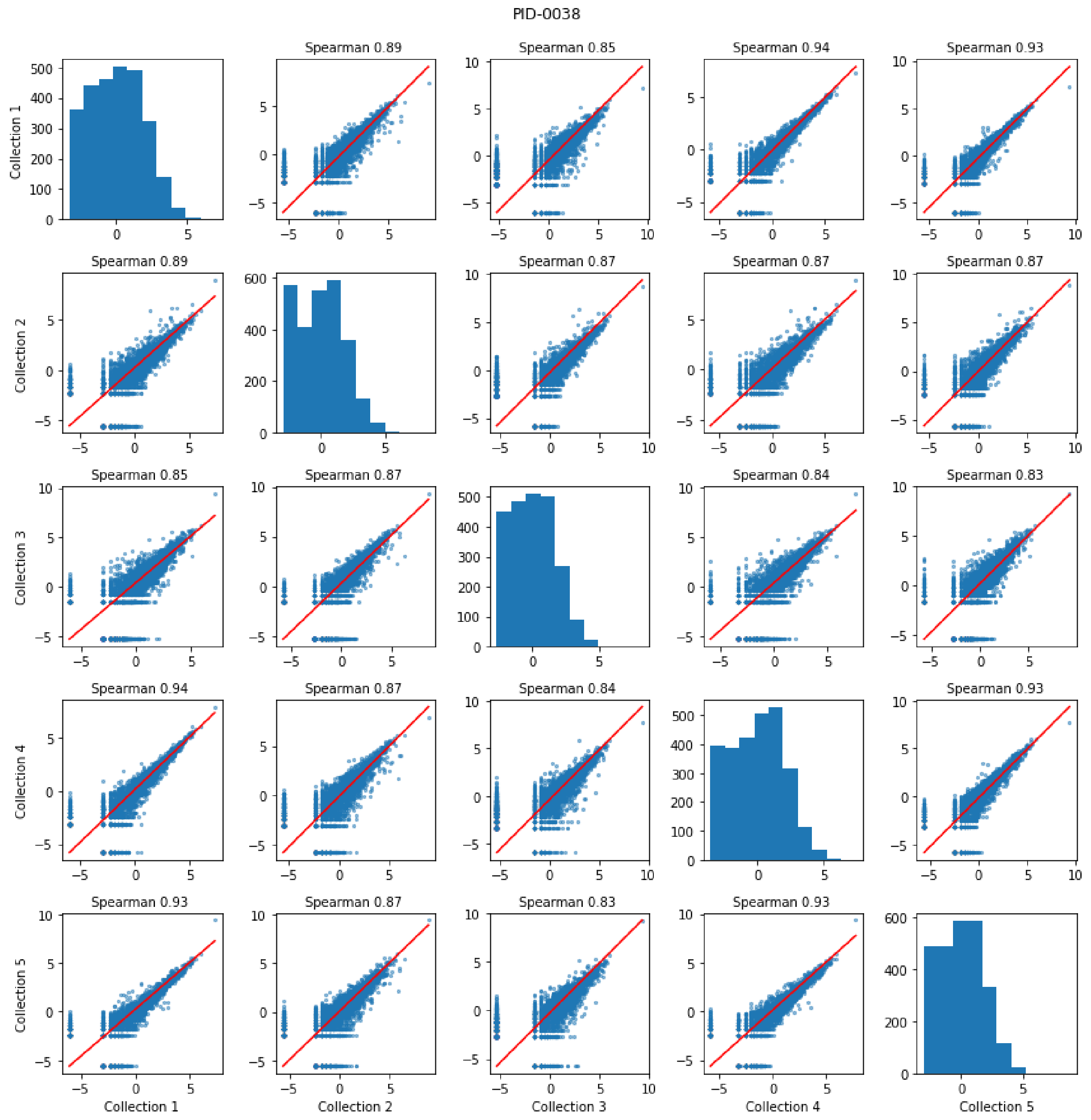
PID-0038
Spearman 0.89
Spearman 0.85
Spearman 0.94
Spearman 0.93
Collection 1
Spearman 0.89
Spearman 0.87
Spearman 0.87
Spearman 0.87
Collection 2
Spearman 0.85
Spearman 0.87
Spearman 0.84
Spearman 0.83
Collection 3
Spearman 0.94
Spearman 0.87
Spearman 0.84
Spearman 0.93
Collection 4
Spearman 0.93
Spearman 0.87
Spearman 0.83
Spearman 0.93
Collection 5
Collection 1
Collection 2
Collection 3
Collection 4
Collection 5

L.

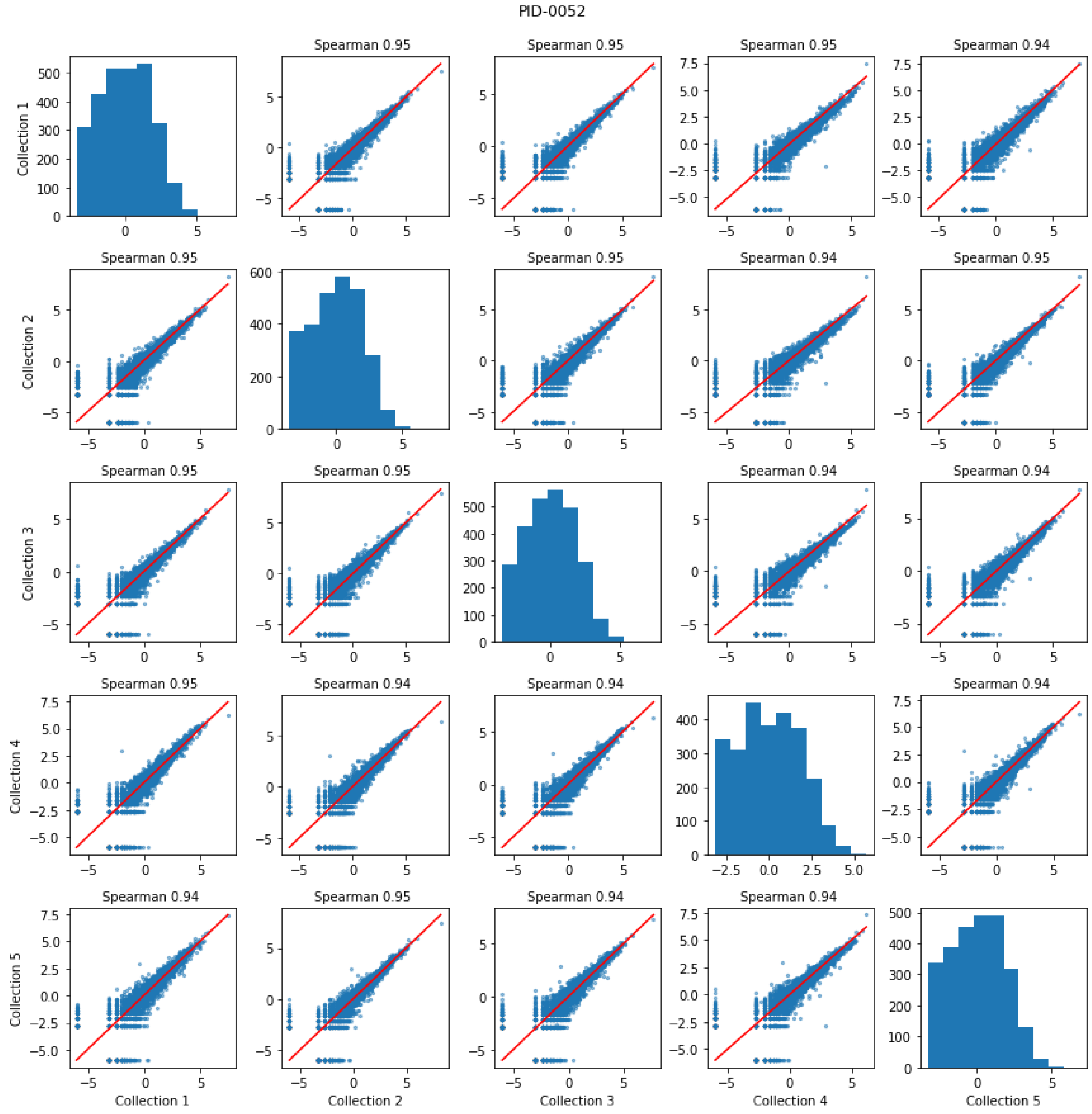
PID-0052
Spearman 0.95
Spearman 0.95
Spearman 0.95
Spearman 0.94
Spearman 0.95
Spearman 0.95
Spearman 0.94
Spearman 0.95
Spearman 0.95
Spearman 0.95
Spearman 0.94
Spearman 0.94
Spearman 0.95
Spearman 0.94
Spearman 0.94
Spearman 0.94
Spearman 0.94
Spearman 0.95
Spearman 0.94
Spearman 0.94
Collection 1
Collection 2
Collection 3
Collection 4
Collection 5

M.

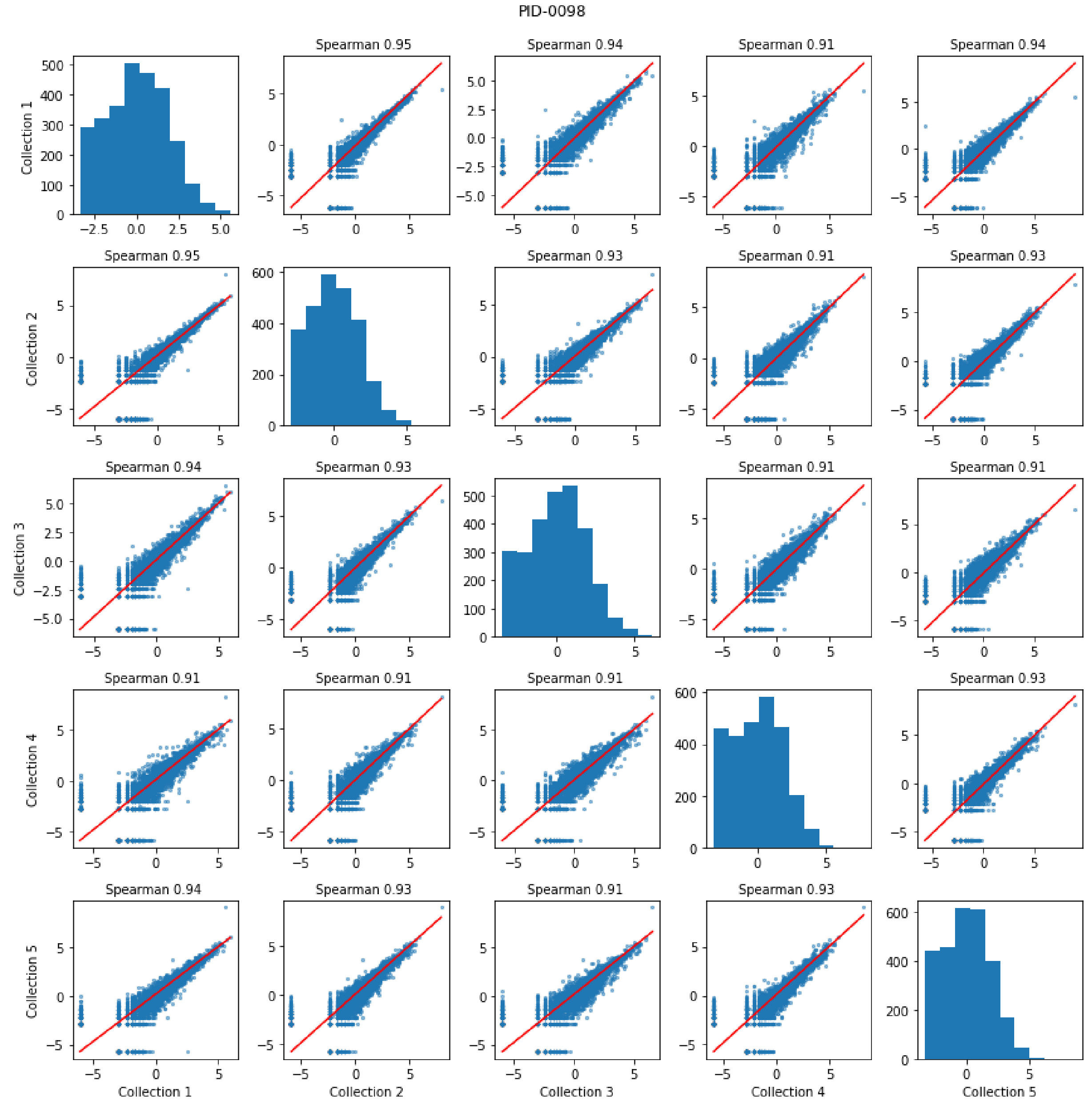
PID-0098
Spearman 0.95
Spearman 0.94
Spearman 0.91
Spearman 0.94
Spearman 0.95
Spearman 0.93
Spearman 0.91
Spearman 0.93
Spearman 0.94
Spearman 0.93
Spearman 0.91
Spearman 0.91
Spearman 0.91
Spearman 0.91
Spearman 0.91
Spearman 0.93
Spearman 0.94
Spearman 0.93
Spearman 0.91
Spearman 0.93
Collection 1
Collection 2
Collection 3
Collection 4
Collection 5

N.

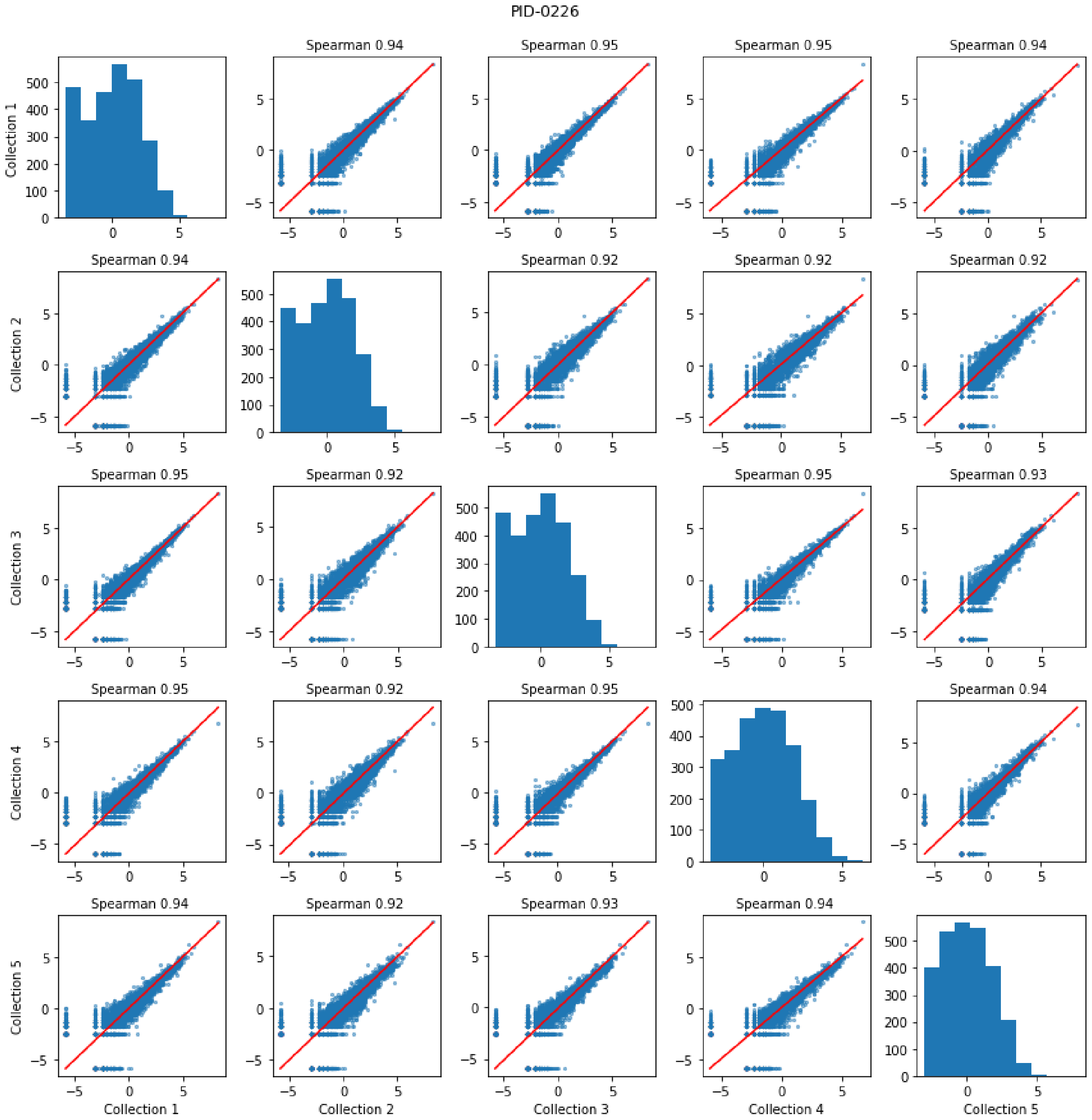
PID-0226
Spearman 0.94
Spearman 0.95
Spearman 0.95
Spearman 0.94
Spearman 0.94
Spearman 0.92
Spearman 0.92
Spearman 0.92
Spearman 0.95
Spearman 0.92
Spearman 0.95
Spearman 0.93
Spearman 0.95
Spearman 0.92
Spearman 0.95
Spearman 0.94
Spearman 0.94
Spearman 0.92
Spearman 0.93
Spearman 0.94
Collection 1
Collection 2
Collection 3
Collection 4
Collection 5

O.

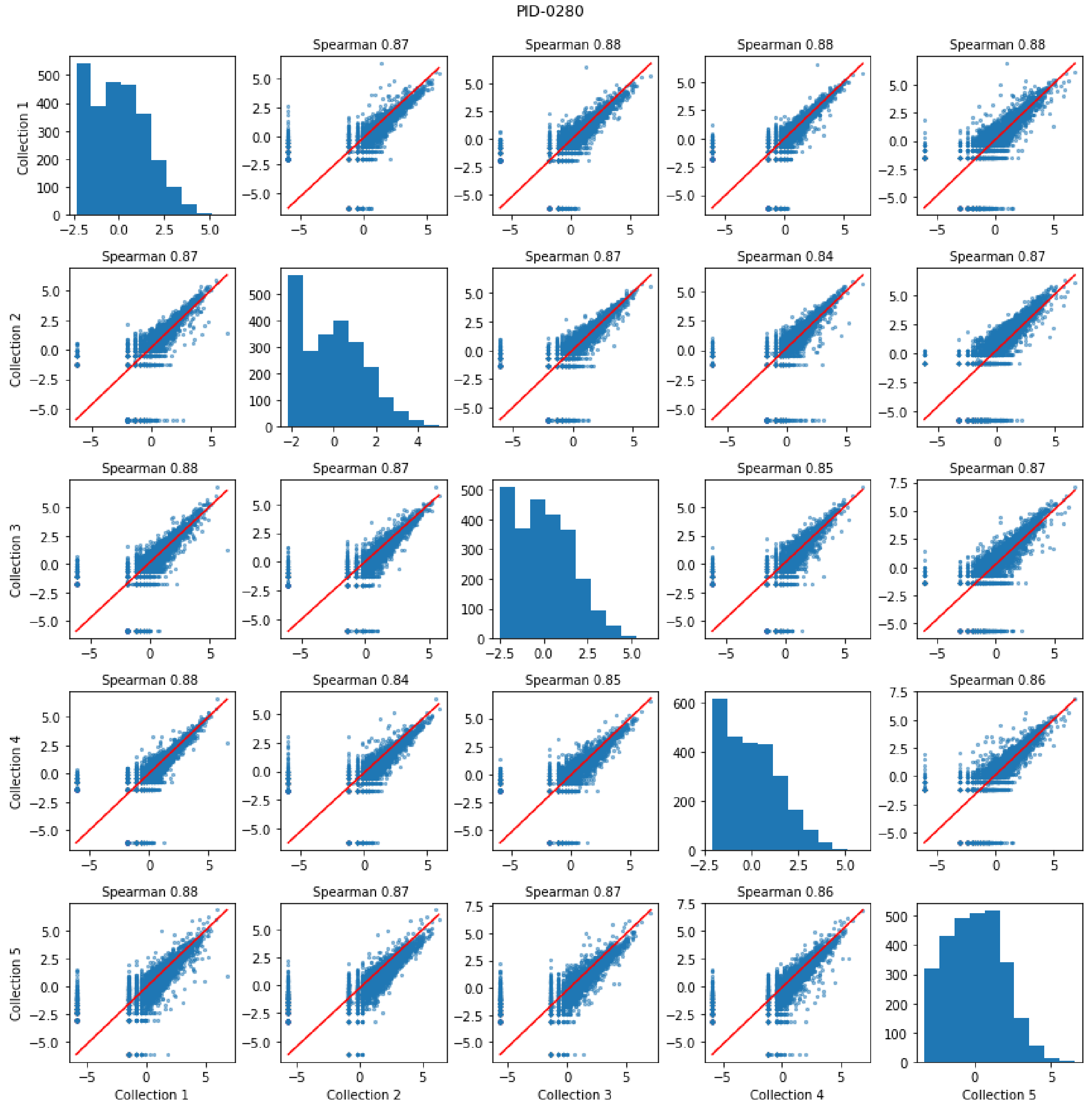
PID-0280
Spearman 0.87
Spearman 0.88
Spearman 0.88
Spearman 0.88
Spearman 0.87
Spearman 0.87
Spearman 0.84
Spearman 0.87
Spearman 0.88
Spearman 0.87
Spearman 0.85
Spearman 0.87
Spearman 0.88
Spearman 0.84
Spearman 0.85
Spearman 0.86
Spearman 0.88
Spearman 0.87
Spearman 0.87
Spearman 0.86
Collection 1
Collection 2
Collection 3
Collection 4
Collection 5

P.

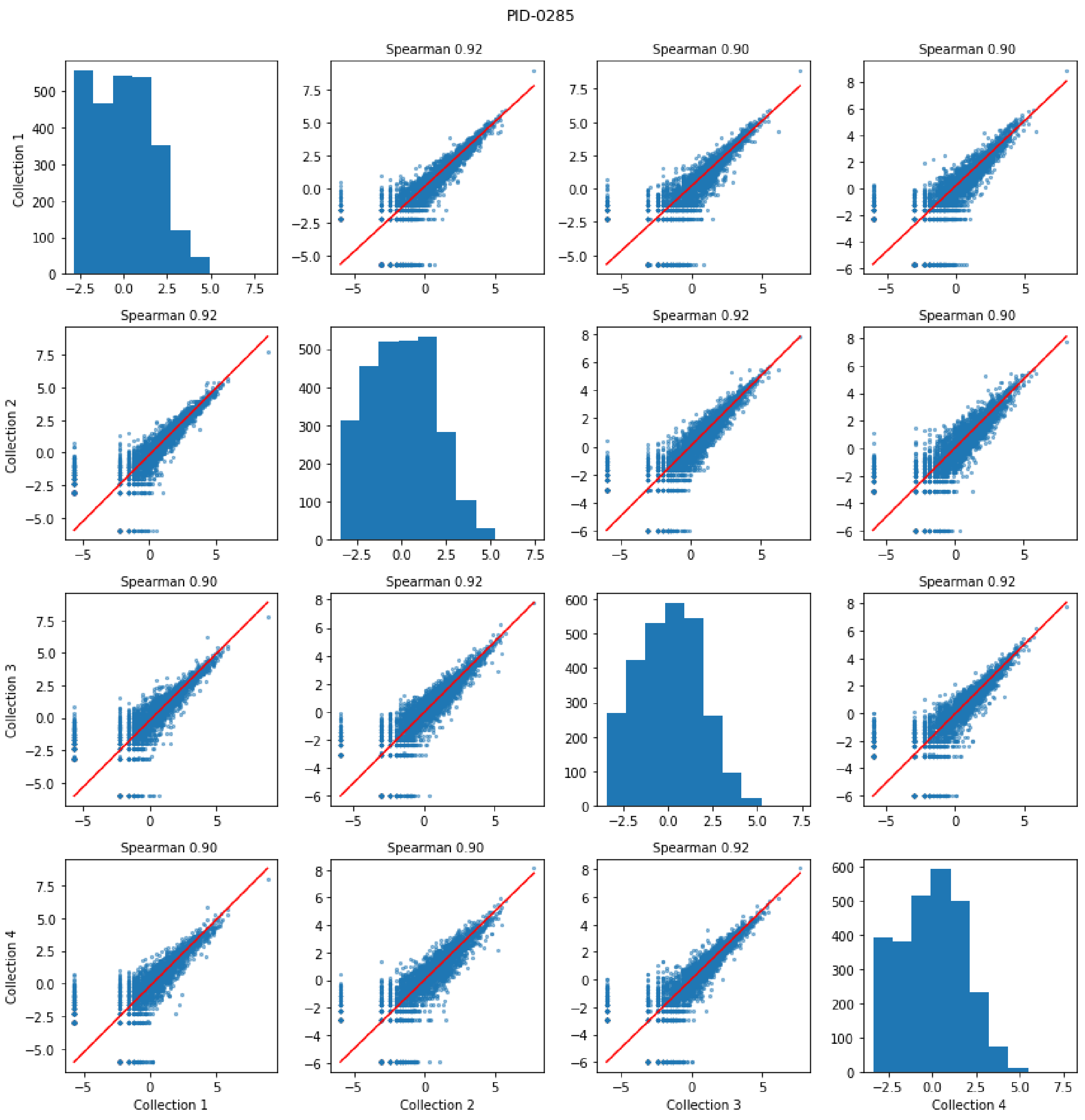


**Supplemental Figure 1.** Longitudinal stability of saliva transcriptome in eight study participants over a 5 week period. (A-H) Scatter plots comparing the relative abundance values of each taxon at each collection time. (I-P) The same analysis was repeated for KOs comparing sum transcripts per million (tpm) of each KO.
